# Evolutionary rescue in toxic combinations of metal is dependent on metal interactions

**DOI:** 10.64898/2026.10.16.759775

**Authors:** Penelope C. Kahn, Sarah P. Otto

**Author notes:** Open Research statement: Data and novel code are provided in an external repository (https://github.com/pennykahn/ComboMetalEE) to be evaluated during the peer review process.

## Abstract

Populations facing abrupt environmental change may avoid extinction through evolutionary rescue, in which beneficial mutations allow individuals to survive and reproduce under otherwise lethal conditions. However, theory on evolutionary rescue has largely focused on single stressors, leaving open how interactions among simultaneous stressors shape adaptation. In particular, synergistic interactions, which amplify toxicity, and antagonistic interactions, which reduce it, may impose distinct selective pressures and lead to different fitness outcomes. To investigate this, we experimentally evolved *Saccharomyces cerevisiae* under lethal combinations of six metals (Cd, Co, Cu, Mn, Ni, and Zn), obtaining rescued lines in eight of the fifteen pairwise combinations. We then measured the fitness of evolved lines in the original metal combination, each component metal individually, and the ancestral environment. Adaptation differed systematically with metal interactions: lines evolved in antagonistic combinations generally gained fitness in the individual metals as well as in the combination, whereas lines evolved in synergistic combinations tended to show more specialized adaptation, with fitness benefits concentrated in the combination and reduced fitness in the component metals. No rescue was observed in combinations whose toxicity was consistent with concentration addition, although differences in the precise toxicity of experimental treatments may have influenced rescue outcomes. These findings demonstrate that interactions among simultaneous stressors can shape not only the likelihood of evolutionary rescue but also the breadth of its fitness benefits, highlighting the importance of accounting for stressor interactions when investigating evolutionary responses to environmental change.

## Introduction

Organisms in the natural world are subject to multiple simultaneous selection pressures, and while we have a well-developed understanding of how evolution occurs in response to individual stressors, our understanding of how selection dynamics change under combined stressors remains limited. Evolve and resequence experiments in recent years have improved our understanding of the molecular basis of adaptation to abiotic stressors, some examples of which include antibiotics Toprak et al. (2012), thermal stress Tenaillon et al. (2012), nutrient limitation Mable (2001), and ethanol stress Voordeckers et al. (2015). Fewer studies incorporate multiple selection pressures into their evolution experiments Phillips et al. (2020); Brennan et al. (2022), and of those that do, even fewer characterize how the interactions among stressors affect evolutionary dynamics.

Given that stressors can interact such that their combined effect deviates from that expected based on the individual impacts (here on growth rate), it remains an open question how those interactions impact the process of adaptation. Synergistic interactions occur when the combined effect of two stressors is more harmful than expected based on their individual effects, whereas antagonistic interactions occur when the combined effect is less harmful than expected. To quantitatively identify these interactions, scientists rely on a null model for non-interaction, such as “concentration addition” (also called Loewe additivity), which assumes that adding equally harmful concentrations of two toxins will have the same effect as doubling the concentration of either one Cedergreen (2014) (described for this system in Kahn and Otto (2025)). Departures from concentration addition are quantified by a, where positive values denote antagonistic interactions, negative values denote synergistic interactions, and the magnitude of a signifies the strength of the interaction (i.e., how much more or less harmful the combination is than expected by concentration addition).

In an applied context, clinical microbiologists and physicians have attempted to leverage synergistic interactions between drugs to improve treatment efficacy (e.g., for microbial infections, HIV, COVID-19, and cancer) by increasing therapeutic potency and decreasing the required dose, thereby minimizing harmful side effects Lehár et al. (2009); Tan et al. (2012); Jin et al. (2021); Duarte and Vale (2022). Some research claims a further benefit to synergistic drug combinations in preventing the evolution of drug resistance Fischbach (2011); Baym et al. (2016). Resistance to combinations of drugs, whether combining synergistically or otherwise, may be harder to evolve because the simultaneous presence of two drugs amplifies the physiological burden on cells. A greater burden may narrow the range of viable mutations and reduce the probability that single-step resistance mutations will be sufficient for survival, thereby increasing the likelihood of extinction before adaptation can occur Michel et al. (2008). Furthermore, once resistance arises, it may have diverse pleiotropic effects: cross-resistance to both drugs, resistance to one stressor over another, or novel resistance mechanisms specific to some emergent property of the interaction Bollenbach et al. (2009); Palmer and Kishony (2013). Understanding the likelihood of generating pleiotropic resistance effects in different drug combinations will be essential for informed clinical practice.

Another relevant scenario where organisms face multiple simultaneous stressors is in environments with high concentrations of heavy metals. Such environments can occur naturally, for example, in serpentine soils and hydrothermal vents, where elevated concentrations of metals like nickel, chromium, and arsenic shape microbial communities Koner et al. (2023); Zhou et al. (2023). While environments with high metal concentrations have been relatively rare for much of Earth’s history, human activities have greatly increased the number of affected habitats. Following industrialization and the technological revolution, humans have induced a mass redistribution of metals for use in countless applications in agriculture and industry Zhou et al. (2020). Destructive bulk extraction methods produce large amounts of waste that must be contained in longterm storage facilities Hudson-Edwards (2016) (i.e., mine tailing ponds), which are vulnerable to failure and release of toxic material into the surrounding environment Rico et al. (2008), with devastating consequences for both humans and wildlife Kossoff et al. (2014); Islam and Murakami (2021). Importantly, these wastes contain mixtures of toxic substances including sulfidic compounds, metal oxides, and multiple soluble metal elements Lottermoser (2010), the components of which can interact, modifying the toxicity of the surrounding environment.

These widespread environmental changes raise the question of how adaptation proceeds when organisms face simultaneous, interacting stressors, particularly in environments where survival depends on rapid evolutionary responses. For example, in the case of evolutionary rescue, a population that has been stable for some time may suddenly encounter a drastic environmental shift that most individuals cannot tolerate and pushes the population toward extinction. Survival of the population then hinges on the chance presence of a beneficial allele – whether already present in standing variation or introduced through mutation or migration – that enables one or a few individuals to persist, reproduce, and ultimately rescue the population. However, when multiple interacting stressors are present, these dynamics may become more complex. For instance, the need to resist two stressors may constrain the possible adaptations available to populations, potentially requiring either pleiotropic mutations or multiple independent beneficial mutations. Pleiotropic effects may arise that enable survival in both components of the stressor combination, or, conversely, interactions between selective pressures could prevent adaptation to one or both, depending on the relative strength of the stressors, their modes of toxicity, and the “fitness landscape” describing their interaction.

Different metals have been shown to interact in many studies Weltje (1998); Verslycke et al. (2003); Horvat et al. (2007); Balistrieri and Mebane (2014); Gao et al. (2020). For example, in Kahn and Otto (2025), I measured growth in 15 pairwise combinations of six metals (Cd, Co, Cu, Mn, Ni, and Zn) and found a variety of interactions ranging from strongly synergistic in combinations with copper to strongly antagonistic in combinations with manganese. The extent to which individual metals dominated interactions with others can be expressed in the following series: Mn_(+)_ *>* Cu_(_*_−_*_)_ *>* Ni_(+)_ *>* Cd_(0)_ = Co_(0)_ = Zn_(0)_, where the subscript refers to whether synergistic (*−*) or antagonistic (+) interactions result, determined in Kahn and Otto (2025)) by each metal’s average interaction value, *a*, when combined with other metals. Mn is the strongest interactor, and mixtures containing this metal are less toxic than expected (antagonistic, positive *a* in all combinations), Cu is second strongest, and mixtures containing this metal are more toxic than expected (synergistic, negative *a* except when paired with Mn), and Ni is antagonistic, though less strongly than manganese (positive *a* except when paired with Cu). Cd, Co, and Zn act neutrally in that their combinations adhere to the null model of concentration addition (*a ≈* 0), except when paired with a strongly interacting metal (e.g., positive *a* when paired with Mn).

To investigate how interactions between pairs of metals affect the outcomes of rapid evolution, we conducted a short-term evolution experiment, exposing an isogenic strain of Saccharomyces cerevisiae to binary combinations of six metals at concentrations that (i) imposed equal stress from each metal and (ii) were just over the lethal threshold for the ancestor when combined, requiring evolutionary rescue via random mutation(s) for growth. We then assayed fitness of the rescued lines under four conditions: the original metal pair, each metal individually, and the ancestral environment lacking excess metal. These comparisons allowed us to assess the pleiotropic consequences of adaptation to combined metal stress. We hypothesized that adaptation to a metal combination would generally confer increased fitness in each single-metal condition. However, we expected that strongly interactive elements, like manganese and copper, might disproportionately influence adaptation, for example, by limiting beneficial mutations to those that resolve the emergent stress resulting from a strong interaction between metals.

## Materials and Methods

### Strains and media

The yeast used to found populations in the evolutionary rescue plates were 80 of the 240 ancestral populations used in Bazzicalupo et al. (2025). These founding populations were colony isolates of the ancestor, OLY077 (itself a colony isolate grown from a single haploid MAT*α* W303 cell). We expect a very small amount of standing genetic variation to be present within each well, as some mutations arose during the propagation from a single cell. With a mutation rate of *∼* 4 *×* 10*^−^*^10^ per nucleotide (or 0.005 per genome) per generation (Sharp et al., 2018), the number of mutations arising during an overnight culture is expected to be small, with most occurring in the last few cell divisions and so being at low frequency (more quantitative description in subsection Evolutionary rescue experiment). As shown in Bazzicalupo et al. (2025) (Supplemental Information), there were no shared mutations among the evolved lines from the same well position in different plates (inoculated from one of the 80 colony isolates), providing empirical evidence that the number of mutations occurring during colony expansion and overnight culture was very low, resulting in nearly isogenic populations both within and between wells. Importantly, because the 80 colony isolates were each propagated from different single cells, any standing genetic variation that arose would be independent of the other colony isolates (not shared variation). We refer to these 80 colony isolates as the “founding populations”.

All liquid media used in the evolutionary rescue experiments and fitness assays contained a base of YPD (yeast extract, peptone, dextrose; RPI Research Products Y20090) + adenine, hereafter referred to as YPAD, to which volumes of aqueous metal solutions or sterile ddH_2_O (controls) were added to achieve desired metal concentrations. Stocks of aqueous metal solutions Stock metal solutions (0.1 M) were prepared using the following metal salts in powder form: Cadmium sulfate, anhydrous, ACS, 99+% (Alfa Aesar 11861); Cobalt(II) sulfate, heptahydrate, 98% (Alfa Aesar A16201); Copper(II) sulfate, pentahydrate, 98+% (Sigma-Aldrich C3036); Manganese(II) chloride, tetrahydrate, 99+% (Sigma-Aldrich M3634); Nickel(II) sulfate, hexahydrate, ACS, 98% (Alfa Aesar 36336); Zinc sulfate, heptahydrate (Sigma-Aldrich Z0251). Measured powder was added to distilled-deionized, ultra-pure water (Barnstead B-pure and Barnstead Nanopure Diamond) and stirred until crystals dissolved. The solutions were autoclaved at 121 ◦C and 15 psi with a sterilization time of 20 minutes.

Growth assays were conducted using the Bioscreen C microbiology workstation (Oy Growth Curves Ab Ltd., 2009), which automates the measurement of microbial growth by recording turbidity of liquid cultures in 100-well plates at set time intervals. Cultures were incubated at 30 *^◦^*C with continuous plate shaking between measurements, and absorbance readings were collected using wide band filters. For all assays, fitness was quantified as the maximum mitotic growth rate in the Bioscreen C (see details below). Additional methodological details and analysis code are available in Gerstein et al. Gerstein et al. (2012) and at https://github.com/joelkcamp/crossTolerance_bazzicalupoEtAl.

We tested for respiratory phenotype by spot plating 5 *µ*L of 1:100 diluted overnight culture on solid petri plates containing YPD (fermentable carbon source) and YPG (yeast extract, peptone, and glycerol, a non-fermentable carbon source). If cells could grow on both media types, they retained respiratory function (”grande” phenotype). If cells only grew on YPD, they had lost respiratory function during evolutionary rescue (”petite” phenotype).

### Evolutionary rescue experiment

For evolutionary rescue in each metal combination, we chose concentrations of metals with two goals: (i) equivalent stress imposed by each metal in isolation and (ii) stress when combined that just prevented detectable growth. Here, “stress” refers to reduction in maximum growth rate (*r*), with a lower *r* indicating a higher stressor intensity. A typical growth curve of the ancestor in YPAD yields an *r* of around *∼*0.3, and *r* calculated from typical noise in a blank sample (lacking yeast) yields an *r* of *∼*0.005*−*0.045 (Figure S2 in Kahn and Otto (2025)). To find the lowest metal concentrations that met these criteria, we performed checker-board assays Bellio et al. (2021), which use a matrix of concentration combinations to assess interactions between two stressors (initially developed for antibiotics). Detailed methods can be found in the Supplementary Methods, examples of checkerboard assay results can be found in Figure S1, and concentrations chosen for evolutionary rescue can be found in Table S1.

The evolution environments were prepared in 96 deep-well plates, one plate per combination. A batch of metal media was prepared by mixing volumes of 0.1 M aqueous metal solutions (0.001 M for Cd) into sterile liquid YPAD for a final volume of 150 mL (concentrations in Table S1). After thorough mixing, the media was poured into a sterile reservoir. 1 mL of media was transferred to each well of the evolution environment plates using an 8-lane multi-channel pipet. Plates were then sealed with silicone mat lids to prevent evaporation and contamination while founding populations were prepared for inoculation. Leftover media was poured into 50 mL Falcon tubes and stored at room temperature for later use in confirming growth of putative evolved populations in metal media.

Cultures of founding populations were prepared the day before by introducing a small frozen sample, scraped by pipet tips of a multichannel pipet, into the wells of a 96 deep-well plate containing 1 mL YPAD per well, and shaking overnight at 30 *^◦^*C. As described above, we expect a small number of mutations to arise during propagation (*∼*28 doublings based on final cell density estimation in Table S2; *∼*0.005 mutations per genome per cell generation Sharp et al. (2018); *∼*0.14 mutations per cell across all doublings; *∼*18,000 cells founding each well, Table S2). Rescue mutations may have been present in the initial genetic variation within each founding cell population or may have arisen *de novo* once in the metal environment, but the relative contribution of these two sources is unknown. Before inoculating founding populations into the evolution environments, samples of overnight cultures were measured for OD, and all cultures were diluted down to an OD of 0.2, then further diluted 1:10, and 10 *µ*L was added to 1 mL metal media in the corresponding middle 80 wells of the evolution environment plate, leaving the outer 16 wells blank to monitor for cross-contamination between wells. The mat lids of the evolution environment plates were resealed, and the plates were incubated at 30 *^◦^*C, shaking at 250 rpm, for 14 days.

Each day after inoculation, all wells in the evolution environment plates were inspected for visible growth. The plates were removed from the incubator and left stationary for ten minutes to allow growth to settle to the bottom of the wells. A 96 LED array lamp the same size as the plates was held to the lid and wells were visually inspected from the bottom. Visible growth appeared as a dark mass when illuminated from above and a white mass with defined edges when illuminated from below. As the onset of visible growth was often very faint and could be confused with a collection of cellular debris or media precipitate, the diameter of the mass was measured, and qualitative observations were recorded. If the size of the mass increased over the next 24 hours, the well was sampled. The typical mass size at time of sampling was *∼*1-2 mm in diameter, about the size of a colony.

For sampling, a well containing a putative evolved population (visible mass) was carefully and thoroughly mixed by pipetting up and down. To prepare a freezer stock, 400 *µ*L was transferred to a cryotube containing 400 *µ*L 30% glycerol and stored at *−*80 *^◦^*C (referred to as “original sample”). Metal resistance of the putative evolved population was confirmed by testing for growth in 2 mL metal media: 200 *µ*L was added to a glass tube containing 2 mL leftover metal media from preparation of the evolution environments. If OD of the culture surpassed 0.5 within four days of incubation on a rotator at 30 *^◦^*C, metal resistance of the evolved population was confirmed. Any cultures that did not display growth were eliminated from further experimentation. Contamination of a blank well by yeast occurred twice: in well H1 of the CdNi plate and H12 of the CoCu plate; there was no contamination by any other species. This indicates that cross-contamination between wells is possible albeit unlikely. Whole-genome sequencing of these lines (not included in this thesis) will reveal the true extent of cross-contamination through identical mutations shared between wells.

While some metal combinations yielded evolved populations, others did not, so a second trial of the evolution experiment was performed for the combinations without any rescued populations, raising the metal concentrations if growth was observed in all wells in the first trial and lowering them if growth was not observed in any wells. Across both trials, eight combinations displayed growth in some, but not all wells. Concentrations used in both trials can be found in Table S1.

We isolated a single colony to represent each evolved population. We did not measure genetic diversity within an evolved population, but we expect it to be relatively low given that (i) the extreme selection pressure is expected to favour rare, large-effect mutations, which is supported by the low frequency of wells displaying growth (mean 24% ± SE 4%) and (ii) wells were sampled shortly after growth was detected, minimizing the time a rescued population had to differentiate beyond the first-step mutations. Furthermore, we chose to use colony isolates for fitness assays (and future whole-genome sequencing) to minimize the within-group variation, which will allow us to confidently match genotype to phenotype data in future analyses (not included in this thesis).

Frozen scrapes from the cryotubes containing the original evolved population samples were streaked onto solid YPAD petri plates and incubated at 30 *^◦^*C for two days to generate colonies from single cells. An average-sized colony was picked with a toothpick, inoculated into 2 mL liquid YPAD, and grown overnight in a rotator at 30 *^◦^*C. If there were large and small colonies from the same population, which was the case for about 15% of the populations, an average-sized colony from the more frequent type was chosen. As evolved yeast often had decreased growth rates compared to the ancestor, these cultures were grown without excess metal to maximize cell density in the overnight culture at the time of sampling for freezer stocks. The next day, these cultures were sampled for *−*80 *^◦^*C freezer stocks using the previously described method to obtain isogenic representatives that we refer to as “evolved lines”. Detailed data from the evolution experiment can be found in Table S2.

All experiments and future sequencing is performed on the evolved lines rather than whole populations, with the exception of respiratory phenotype testing. Testing growth of both samples on fermentable and non-fermentable media allowed us to observe the prevalence of heterogeneity within the evolved populations (sampled directly from deep-well evolution plates) with respect to petites and grandes. For example, if the evolved population could grow on both YPD and YPG, but the evolved line representative could only grow on YPD, this indicated we chose a petite colony from a mixed population containing both petites and grandes.

### Fitness assays

For each metal combination, we measured the fitnesses of evolved lines and the ancestor in four test environments: the metal combination, each metal individually (at the same concentrations as in the combination environment), and the ancestral environment (YPAD). The assays were broken up into seven batches (Table S4), measuring growth rates of yeast lines evolved in a single metal combination in each batch (except the third batch which assayed both CdMn and NiZn lines). The first six batches were performed over a period of 34 days using the same stock of YPAD media and aqueous metal solutions; the last one was performed 100 days later using new YPAD media but the same metal solutions as the first six batches. Batch effects from conducting these measures on different days can be observed in the slightly variable growth rates of the ancestor in YPAD (grey line at top of graphs in Figure 2A; Table S4).

**Figure 1:**
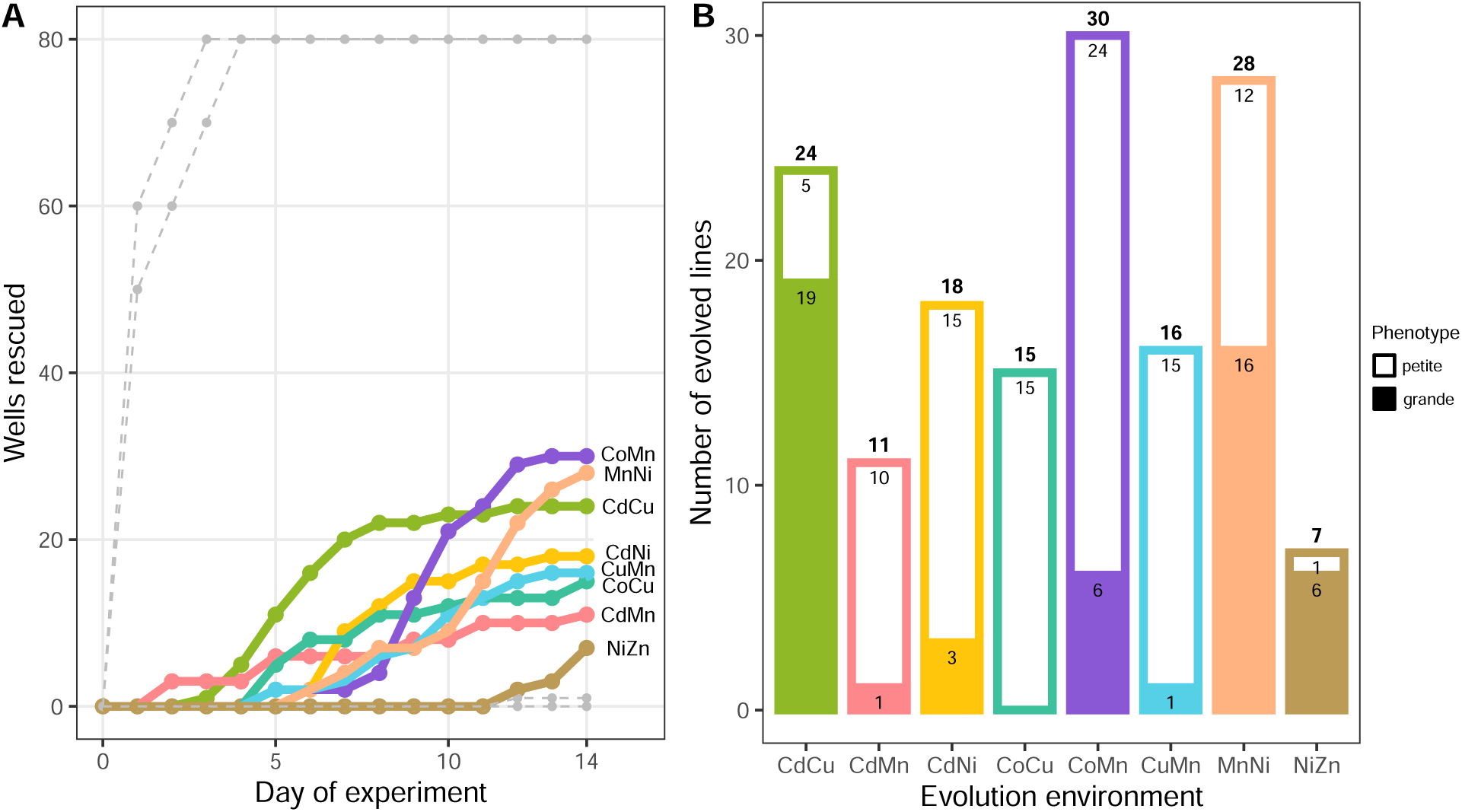
Yeast lines generated during the evolutionary rescue experiment. (A) Time series of wells displaying visible growth over 14 days. Replicate founding populations were inoculated at low density into the middle 80 wells of one 96 deep-well plate for each metal combination. The combinations with rescued lines are displayed in colour. For some combinations, shown in grey, the chosen metal concentrations allowed growth in all or none of the 80 wells (see Table S1) and were thus excluded from further experimentation. (B) Number of evolved lines from each metal combination with breakdown of phenotype by respiratory function (petite is loss of function, grande is wild-type function). Total number of evolved lines from each combination is displayed above each bar.

**Figure 2:**
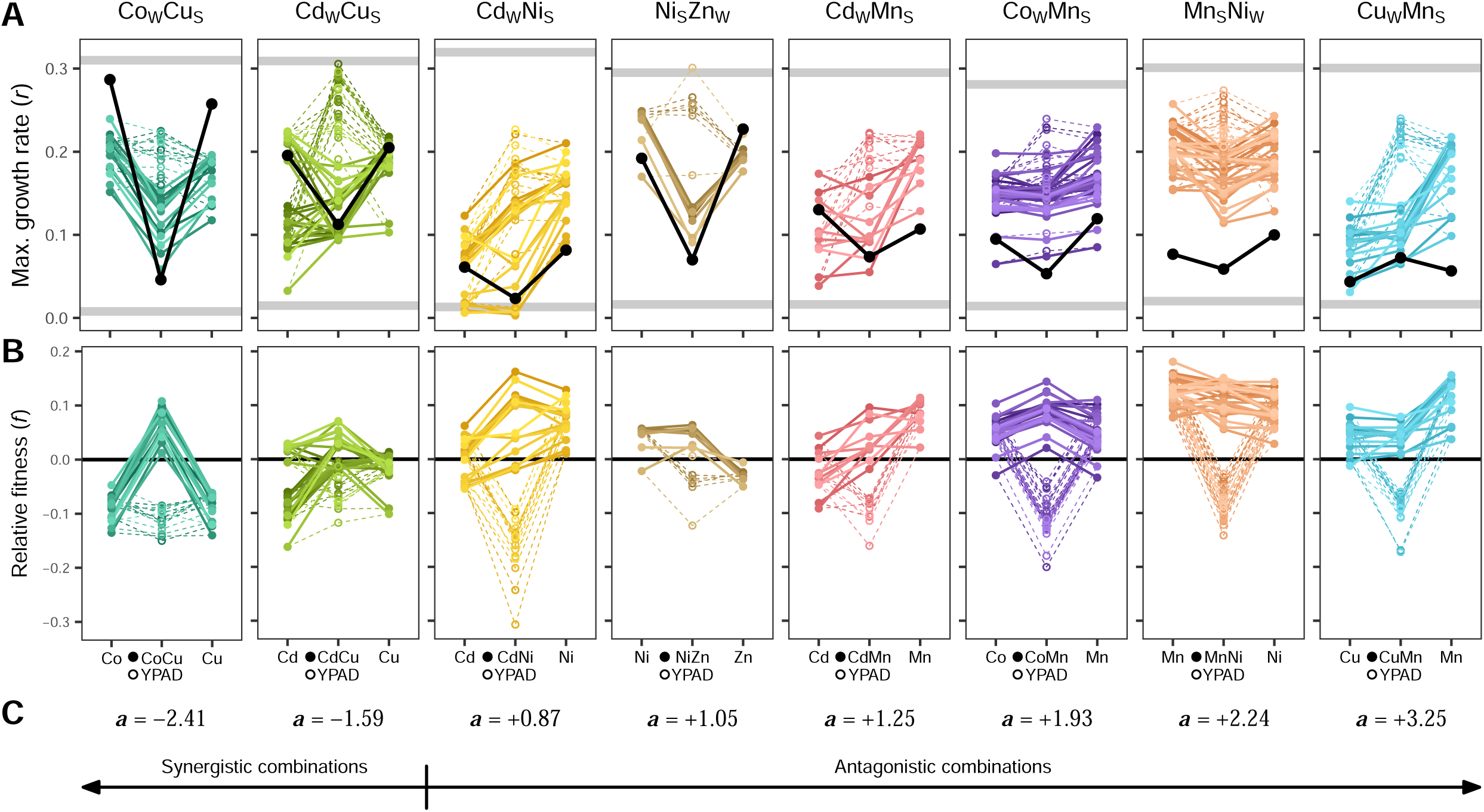
Fitness of all evolved lines in four test environments (metal combination, YPAD, and each single metal). (A) Maximum growth rates (*r*; y-axis) of evolved lines (in color) and the ancestor (in black) are displayed in four test environments. The metal combination in which yeast lines evolved is given above each graph. The grey bars at top and bottom represent 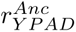 and 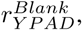 respectively. Growth rates in the single metals are displayed on the left and right sides of each graph; *r_Combo_* and 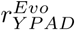 are displayed in the middle as solid and open circles, respectively. Each point represents an average from three technical replicates. (B) Relative fitness (*f* = *r^Evo^ − r^Anc^*; y-axis) of evolved lines (in color) relative to the ancestor (horizontal black line). (C) Interaction value of each metal combination (*a*; deviation from concentration addition) and placement along spectrum of synergism to antagonism as obtained from a previous experiment (see Kahn and Otto (2025)).

The concentrations of metal used to measure fitness of the evolved lines in the Bioscreen C were chosen with similar goals as for the evolutionary rescue experiment: (i) equivalent ancestral maximum growth rates in each single metal, (ii) but instead of aiming for no detectable growth in the combination, we aimed for an ancestral maximum growth rate of 0.05 in the combination. A maximum growth rate of 0.05 is very low but still detectable and was chosen to be similar in harshness to the evolution environments but permissive enough that we could be confident that stressor intensity was standardized across metal combinations.

To choose concentrations that would meet these criteria, we analyzed data from experiments in Kahn and Otto (2025), which measured growth rates of the ancestor (OLY077) in individual metals and their pairs at various concentrations ranging from no growth to normal growth to produce concentration-response curves. The data in Figure 3A of this paper pointed to concentration of the metal combinations that would result in maximum growth rates of *∼*0.05 when combined, while displaying similar growth rates in each of the single metals. After choosing these concentrations, we conducted a preliminary assay testing only the ancestor for confirmation of the growth rates in the single and combined metals. Small adjustments were made to these concentrations based on the preliminary assay results, resulting in the concentrations used in the fitness assays (given in Table S1B).

**Figure 3:**
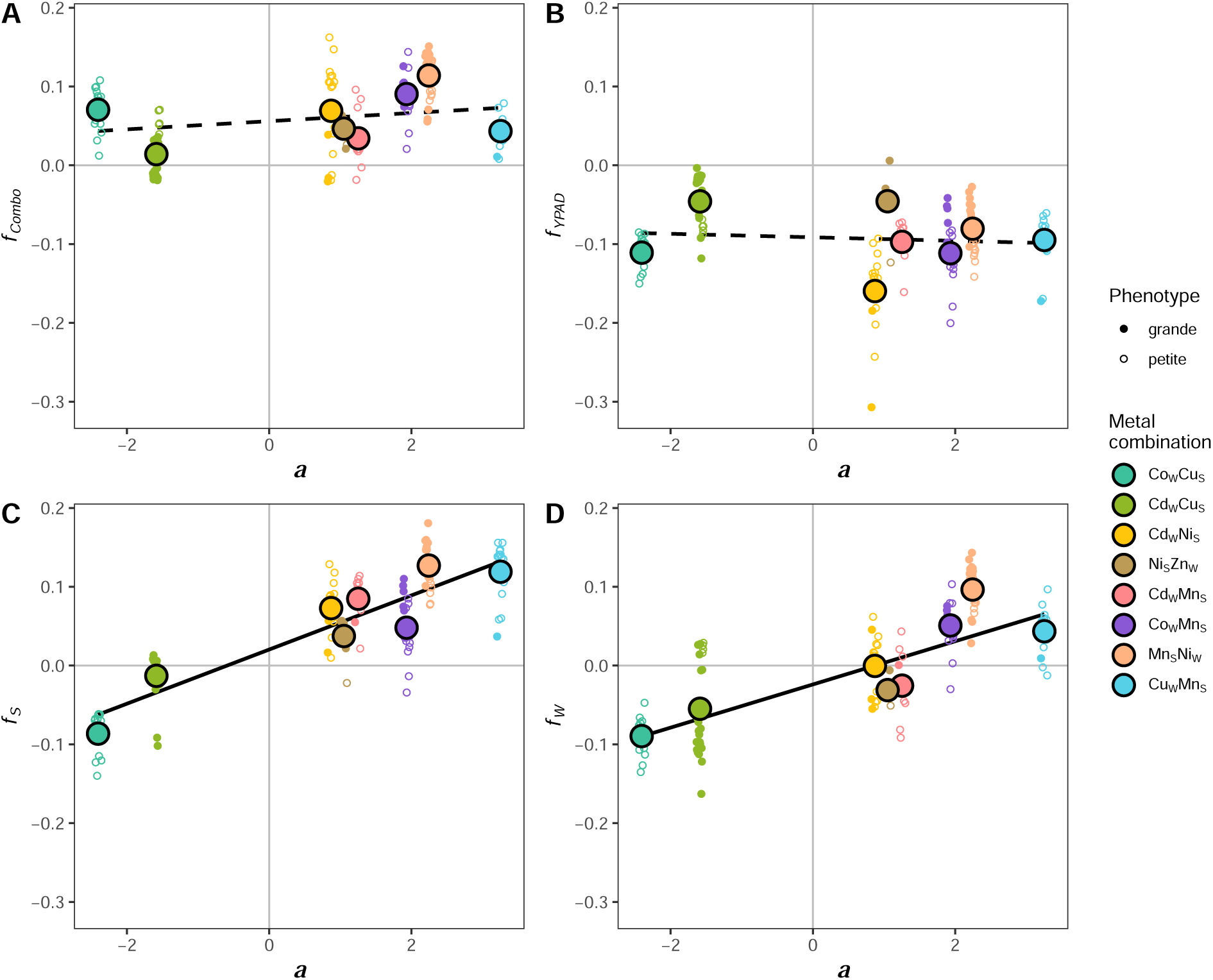
Relationship between interaction value (*a*) and relative fitness in each test environment. All graphs show the following: average relative fitness for evolved lines from metal combinations as large colored dots with black outline; relative fitness of individual evolved lines as small colored dots, open circles representing petites and closed circles representing grandes; linear regressions in black, dashed lines representing non-significant relationships and solid lines representing significant relationships. (A) Relationship between *a* and relative fitness in the metal combination (evolution environment); not significant. (B) Relationship between *a* and relative fitness in YPAD (ancestral environment); not significant. (C) Relationship between *a* and relative fitness in the stronger interactor (slope = 0.034, *R*^2^ = 0.84, *F* (1,6) = 37.12, *p <* 0.001). (D) Relationship between *a* and relative fitness in the weaker interactor (slope = 0.027, *R*^2^ = 0.68, *F* (1,6) = 16.15, *p <* 0.01).

To prepare cultures of evolved lines for use in the fitness assays, frozen scrapes of the cultured colony isolates were inoculated into 2 mL media with metal concentrations at 95% those chosen for the fitness assays and incubated on a rotator at 30 *^◦^*C (for reacclimation to metal combinations following growth in YPAD from colony propagation and subsequent freezing at *−*80 *^◦^*C). Because many of the evolved lines grew slowly compared to the ancestor, their cultures were started three days before the experiment, and the ancestor culture in YPAD was started one day before. If an evolved line grew faster than the others at this stage, a new culture was started from the freezer stock at the same time as the ancestor.

The fitness assays were conducted using the Bioscreen C. Media containing the metal combination and each single metal were mixed in 15 mL Falcon tubes at the concentrations necessary to achieve the targets in Table S1. 480 *µ*L, was aliquoted into 1.5 mL microcentrifuge tubes, one tube per yeast line and one for the blank negative control. The cultures of isogenic samples were measured for OD in a BioTek spectrophotometer and diluted down to an OD of 0.7, then further diluted 1:10 in fresh YPAD. 40 *µ*L of diluted yeast was added to the corresponding 1.5 mL microcentrifuge tubes (40 *µ*L of sterile ddH_2_O was added to the blank negative control tubes). After vortexing, wells were loaded with 120 *µ*L from the 1.5 mL microcentrifuge tubes, such that each well contained *∼*111 *µ*L of media and *∼*9 *µ*L of diluted yeast (approximately 100,000 cells founding each well) arranged to have three technical replicates per treatment randomly arranged across four Bioscreen plates.

Four 100-well Bioscreen C plates were run in parallel at 30 *^◦^*C, shaking at maximum amplitude between automatic OD measurements taken every 15 minutes over 48 hours. Maximum growth rates per hour were determined by calculating the slope of a loess-smoothed growth curve based on the log-OD for each well (using loess in R with degree=1 and span=0.1). When reporting maximum growth rates, the identity of yeast as either evolved or ancestor is given as a superscript, and the identity of the test environment is given as a subscript (e.g., 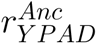 for the ancestor in YPAD and 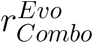 an evolved line in the metal combination).

Relative growth rate (*f* = *r^Evo^ − r^Anc^*), hereafter referred to as “relative fitness”, will always describe evolved lines, so the superscript is blank, leaving only a subscript referring to the test environment (e.g., *f_Combo_* refers to an evolved line’s fitness in the metal combination). For the single metals, we refer to the relative growth rates as *f_S_* and *f_W_*for the stronger and weaker interacting metal from the series determined in Kahn and Otto (2025) (Mn_(+)_ *>* Cu_(_*_−_*_)_ *>* Ni_(+)_ *>* Cd_(0)_ = Co_(0)_ = Zn_(0)_). There were no evolved lines generated in combinations with approximately equal interaction strengths (CdCo, CdZn, CoZn). The relative fitnesses offer a normalized view of changes in *r* across test environments (e.g., the metal pair and the stronger interactor from a given combination) and allow for comparison of changes in *r* across combinations. For comparisons across metal combinations, all data for a combination were averaged and treated as a single point to avoid pseudo-replication. Batch effects may have contributed additional noise to these point estimates, reducing the power of these comparisons.

## Results

### Evolutionary rescue

In the first trial, with 80 low-density populations in a 96 deep-well plate for each metal combination, evolutionary rescue occurred in six of the fifteen metal combinations. Growth was observed in 24 out of 80 wells in CdCu, 20 in CdNi, 30 in CoMn, 17 in CuMn, 28 in MnNi, and 7 in NiZn. The rest displayed growth in either all or none of the eighty replicate wells. The differences in proportion of rescued wells in this first trial might have arisen by chance or have been driven by subtle differences in the toxicity of the metal combinations, which was challenging to control precisely (the target concentration being just beyond where the intrinsic growth rate of the lines fell below zero). When the chosen metal concentrations were not sufficiently toxic, growth occurred in all wells (CdCo, CdMn, CdZn, and CoNi). When too toxic, no wells exhibited growth (CoCu, CoZn, CuNi, CuZn, and MnZn), which might be because the environments were too harsh (intrinsic growth rates too negative) and/or because no single-step mutations would allow growth under these experimental conditions.

To generate evolved lines in more metal pairs, we conducted a second trial with the nine pairs that did not generate evolved populations in the first trial, adjusting the metal concentrations based on the previous results (e.g., decreasing concentrations in combinations that displayed no growth; see Table S1). The second trial resulted in successful evolutionary rescue in two additional combinations: 11 evolved populations in CdMn and 17 in CoCu. Again, growth was observed in all wells for CdCo, while now no growth was observed in CdZn and CoNi (which all grew before), nor in CoZn, CuNi, CuZn, and MnZn (none of which had growth in either trial). While it remains possible that the concentrations of these latter four metal combinations were still too harsh (intrinsic growth rates too low), single-step rescue may not be possible when these metals are combined, even when metal concentrations are just slightly beyond where growth can occur. Figure 1A displays the accumulation of evolved populations over the 14-day experiment (see Table S3 for well location and date of rescue for each evolved line).

The evolved lines often exhibited a petite phenotype (loss of respiratory function; Figure 1B and Table S3). Since yeast can produce energy through both respiration and fermentation, cells that have lost mitochondrial function remain viable but typically suffer reduced growth rates compared to their wild-type counter-parts Vowinckel et al. (2021). Petites have been estimated to arise naturally in yeast culture at a rate of 0.1-1%, though this is highly dependent on the strain Day (2013). Since ROS are endogenously produced by the mitochondria during oxidative phosphorylation, loss of mitochondrial function results in a reduction of cellular superoxide levels Vowinckel et al. (2021) and therefore may be a selective advantage in highly oxidative metal environments. The frequency of petites was significantly different across metal combinations (*χ*^2^(7, N = 149) = 57.4, *p <* 0.001, calculated from 2000 permutations because some entries have few counts), with petites observed in the majority of lines for CdMn (10 of 11), CdNi (15 of 18), CoCu (15 of 15), CoMn (24 of 30), and CuMn (15 of 16) but in less than half of the lines for CdCu (5 of 24), MnNi (12 of 28), and NiZn (1 of 7). These results may point to differences in the level of harm to the mitochondria or in the oxidative burden produced by different combinations.

Having also tested the respiratory phenotypes of the whole rescued populations (original samples from evolutionary rescue plates), we found that 23 of 149 evolved populations showed evidence of heterogeneity in respiratory phenotype with co-existence of petites and grandes (1 of 11 CdMn lines, 5 of 18 CdNi lines, 1 of 15 CoCu lines, 4 of 30 CoMn lines, 4 of 16 CuMn lines, 7 of 28 MnNi lines, and 1 of 7 NiZn lines). This indicates that some genetic diversity arose during the process of evolutionary rescue, but in choosing to isolate colonies to represent each population, our experimental design sacrifices the characterization of this diversity in favour of clearly characterizing single rescue genotypes in both whole-genome sequencing (not included in this study) and fitness assays.

### Fitness assays: Growth in combined vs. single metals

To assess the costs and benefits of evolutionary rescue in each metal combination, we measured fitness of the evolved lines in four test environments (the metal pair, YPAD, and each individual metal at the same concentrations used in the pair). Maximum growth rates (*r*) of the ancestor and evolved lines are displayed in Figure 2A; those of the ancestor (*r^Anc^*) are displayed in black, and those of the evolved lines *r^Evo^* are in colour. Maximum growth rates of the evolved lines relative to the ancestor (i.e., relative fitness, *f* = *r^Evo^− r^Anc^*) are displayed in Figure 2B; the horizontal black line at f=0 represents the ancestor, and colored points above and below this axis represent higher and lower *r^Evo^* compared to *r^Anc^*, respectively. The graphs in Figure 2A and 2B are arranged left to right according to the interaction value (*a*) of the metal combination from Kahn and Otto (2025), as given in Figure 2C. Combinations whose effects are more than additive (i.e., more toxic than expected) are synergistic (negative *a*; left side of Figure 2); combinations whose effects are less than additive (i.e., less toxic than expected) are antagonistic (positive *a*; right side of Figure 2).

The effort to choose concentrations that would induce (i) a low, but detectable ancestral growth rate in each combination 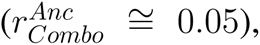 and (ii) equivalent ancestral growth rates in each individual metal 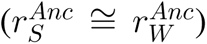 was successful, with an average 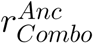 of 0.064 (SE = 0.009) across combinations, and an average absolute difference between 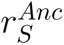 and 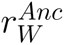 of 0.022 (SE = 0.003). The maximum growth rate of the ancestor in the standard YPAD medium, 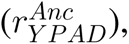 was consistent between batches with an average 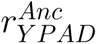 of 0.301, displayed by grey bars at the top of each graph (SE = 0.004); the grey bars at the bottom represent *r* calculated from noise in OD measurements of blank media (no yeast added). Maximum growth rates (Figure 2A) and relative fitnesses (Figure 2B) of the evolved lines in YPAD (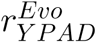 and *f_Y PAD_*) are represented by open circles and are connected to the maximum growth rates and relative fitnesses of the evolved lines in individual metals (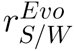 and *f*_S/W_) by dashed lines.

We consider a yeast line’s maximum growth rate in YPAD (*r_Y PAD_*) to be representative of its “intrinsic growth capacity”, its maximum potential for growth in rich media. A greater difference between 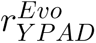 and 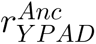 indicates a larger cost of the mutations allowing rescue of an evolved line. Conversely, a greater difference between 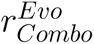 and 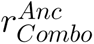 indicates a larger benefit of the underlying mutations.

On average, the magnitude of fitness benefit in the evolution environment resulting from evolutionary rescue (*f_Combo_*) was positive across all evolved lines (Figure 2B) with an average of 0.065 and standard deviation of 0.047, indicating considerable variation. Evolved lines from MnNi and CoMn had the highest growth in the metal combination relative to the ancestor (mean *f_Combo_* [SE]: MnNi, 0.114 [0.005]; CoMn, 0.090 [0.004]), while evolved lines from CdCu had the lowest (CdCu, 0.014 [0.006]). On average, the magnitude of fitness cost in the ancestral environment resulting from evolutionary rescue (*f_Y PAD_*) was negative across all evolved lines with an average of *−*0.096 and standard deviation of 0.049, again indicating considerable variation. Evolved lines from CdNi had the lowest fitness relative to the ancestor in YPAD (mean *f_Y PAD_* [SE]: CdNi, *−*0.161 [0.011]), suggestive of large trade-offs in their adaptations, while evolved lines from CdCu and NiZn had the highest relative fitness, suggesting milder costs (CdCu, *−*0.046 [0.007]; NiZn, *−*0.046 [0.015]).

While averaging data from lines evolved in the same combination avoids pseudoreplication in statistical tests comparing combinations, it obscures some interesting patterns exhibited by individual evolved lines. For example, some evolved lines did not have significant fitness gains compared to the ancestor at the concentrations tested in the fitness assay, indicating that the benefit of these adaptations for metal resistance may be modest or only visible in the deep well plates at the more prohibitive concentrations used in the evolution experiments. Furthermore, significant differences between petites and grandes are lost by averaging their fitnesses, but this topic will be examined further at the end of the next subsection.

### Relationship between interaction of the combination and evolved fitness patterns

We investigated if the ways in which metals interact (*a*) influenced the potential and scope for fitness evolution. The likelihood of evolutionary rescue may be affected if an interaction between stressors changes the shape of the concentration-response surface, creating broader or narrower fitness declines across single and double-metal conditions. Evolutionary rescue generated evolved lines in two of four synergistic combinations (*a <* 0; rescue in CoCu and CdCu; no rescue in CuNi or CuZn), zero of three additive combinations (*a* not significantly different from 0; no rescue in CdCo, CdZn, or CoZn), and six of the eight antagonistic combinations (*a >* 0; rescue in CdNi, NiZn, CdMn, CoMn, MnNi, and CuMn; no rescue in CoNi or MnZn). Evolutionary rescue in this system depends, in part, on choosing metal combination concentrations that lie within a specific range: high enough to prevent growth of the ancestor but as low as possible beyond that. Despite our best efforts, the concentrations that failed to rescue may simply have been outside this range. Alternatively, it is possible that, for some combinations, evolutionary rescue simply may not have been possible in the context of this experiment. We explore this further in the discussion and focus now on the combinations that produced evolved lines.

Next, we examined associations between the interaction value (*a*) and fitness in the four test environments. Such associations provide insights into how adaptive mutations in combinations with different interaction types change the concentration-response surface (difference in shape of colored and black lines in Figure 2A). Figure 3 examines how the interaction value (*a*) of a metal pair relates to the fitness consequences of rescue in each test environment. We did not observe that the interaction between metals in a combination (*a*) influenced the fitness benefits in the metal combination (*f_Combo_*) or costs in YPAD (*f_Y PAD_*) resulting from evolutionary rescue (Figure 3A and 4.3B, respectively). By contrast, *a* was significantly correlated with relative fitness in both single metals, *f_S_* and *f_W_* (Figure 3C and 4.3D); in more antagonistic combinations where metal combinations are less toxic than expected (*a >* 0, right side of the graphs), adaptation to the combination tends to benefit growth in the individual metals. In synergistic metal combinations that were more toxic than expected (*a <* 0, left side of the graphs), however, many of the lines that grew better in the combined metals grew worse in the single metals (CoCu, CdCu; Figure 2A). This relationship may reflect emergent harmful properties from combining these metals that provide a specific selection pressure differing from those imposed by each isolated metal. The relationship between a and relative fitness is slightly stronger in the stronger interactor (*f_S_ ∼ a*: slope = 0.034, *R*^2^ = 0.84, *F* (1,6) = 37.12, *p <* 0.001), compared to the weaker interactor (*f_W_ ∼ a*: slope = 0.027, *R*^2^ = 0.68, *F* (1,6) = 16.15, *p <* 0.01), suggesting that evolution in combinations with a strongly interacting metal (Mn, Cu, Ni) led to greater fitness changes in that single metal compared to the other metal in the pair.

Figure 4 explores how fitness gains in the individual metals trade off against one another among evolved lines within each metal combination. By plotting relative fitness in the weaker interactor (*f_W_*) against relative fitness in the stronger interactor (*f_S_*), we can assess whether adaptations tend to favor one metal at the expense of the other, or whether some lines achieve balanced improvements across both. For most metal combinations, the rescued lines had higher fitness in the stronger interactor than the weaker interactor, falling below the 1:1 line in Figure 4A. Interestingly, the only two groups of evolved lines that do not fit this pattern both evolved in metal combinations containing cobalt. Specifically, lines evolved in CoCu and CoMn tended to have approximately equal fitness in each single metal. Evolved lines from some environments (e.g., CdNi and MnNi) have different fitnesses in the individual metals (points below 1:1 line in Figure 4A), and the magnitude of difference is similar across replicate lines in each group (slope near 1; i.e., individual points have similar distance from 1:1 line). This can be compared to evolved lines from other environments (e.g., CdMn and CuMn) where the magnitude of the fitness difference in individual metals is more variable across. This may point to more diverse mechanisms of adaptation in the latter group compared to the former, leading to more diversity in the pleiotropic effects for fitness in the individual metals.

**Figure 4:**
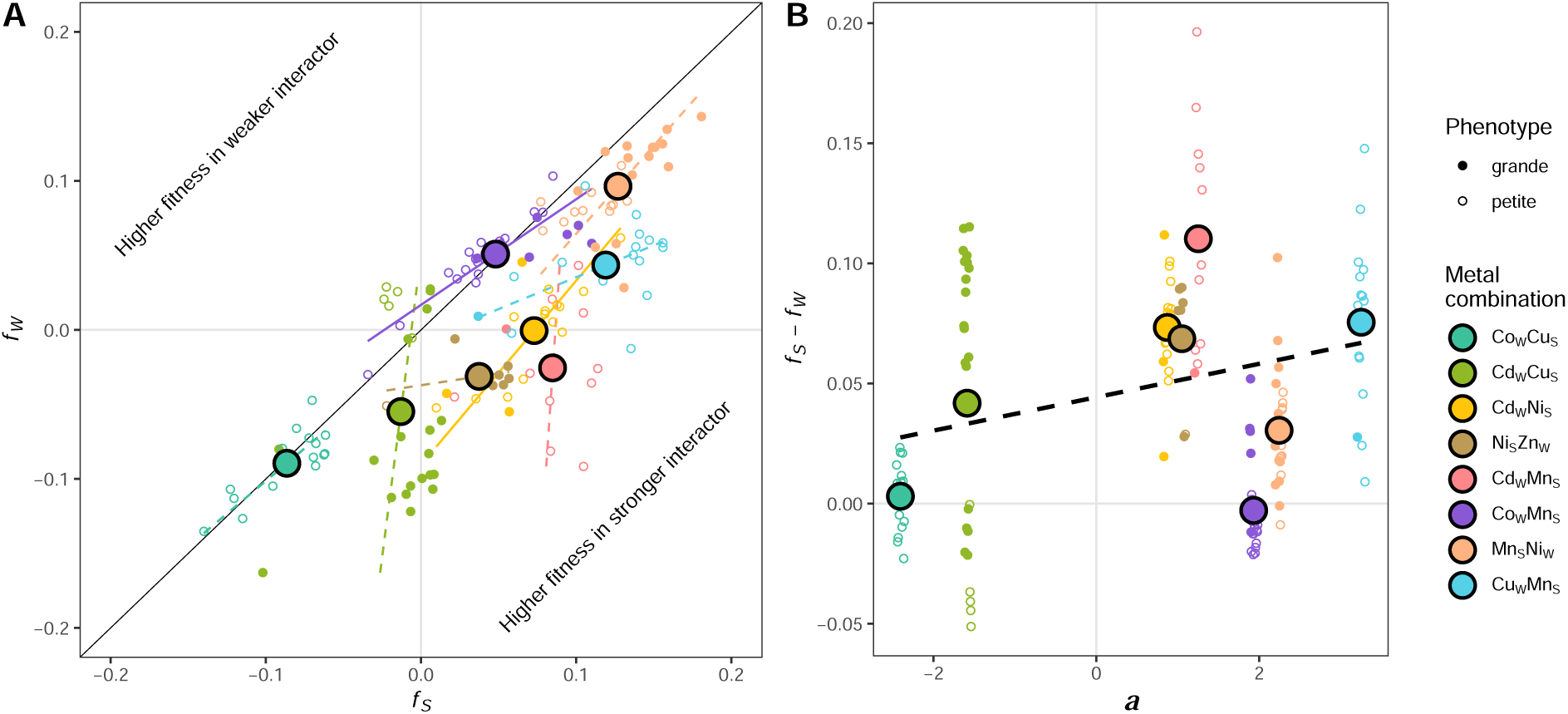
Difference between relative fitnesses in individual metals. Both graphs show average relative fitness for evolved lines from metal combinations as large colored dots with black outline, relative fitness of individual evolved lines as small colored dots (open circles representing petites and closed circles representing grandes). (A) Relationship between relative fitnesses in the stronger interactor (*f_S_*; x-axis) and weaker interactor (*f_W_* ; y-axis). Identities of the stronger and weaker interactors for each combination are shown as subscripts in the legend. 1:1 line is shown in black. Points to the bottom-right of the 1:1 line have higher relative fitness in the stronger interactor, and points to the top-left of the 1:1 line have higher relative fitness in the weaker interactor. Coloured lines show results from Deming regressions, a method to obtain linear fits to data when both axes are measured with error (solid: significant for CdNi and CoMn; dashed: non-significant) indicate significant slopes. (B) Relationship between interaction value (*a*) and difference between relative fitness in individual metals (*f_S_ − f_W_*). Linear regression shown by black dashed line; slope, not significant; intercept, *p* = 0.03.

We next tested if the relative improvement in the strongly interacting versus weakly interacting metal depends on the interaction term a. For example, a more strongly antagonistic metal (e.g., Mn) may protect cells from harm caused by the second metal, reducing selection pressures from that second metal. Conversely, a strongly synergistic interactor (e.g., Cu) may increase the harm caused by the second metal, thereby requiring adaptation to both. There was, however, no significant relationship between a and the difference in relative fitness between the stronger and weaker interactor (Figure 4B). While it is true that lines tend to improve growth more in the stronger interactor than the weaker interactor (significant positive intercept, *p* = 0.03), the magnitude of this difference does not significantly depend on the degree of antagonism or synergism of the combination (non-significant slope).

To statistically test for differences in relative performance between the combination environment and each individual metal we performed two paired t-tests (Table 1), one comparing *f_S_* to *f_Combo_*, and one comparing *f_W_* to *f_Combo_*. Taken together the results reveal some common patterns among combinations. For lines evolved in CdMn, CdNi, and MnNi, the increase in growth rate was highest in the stronger interactor, intermediate in the combination, and lowest in the weaker interactor (*f_S_ > f_Combo_ > f_W_*), indicating greater adaptation to the stronger interactor. Lines in CuMn were similar, but without a significantly lower growth in the weaker interactor (*f_S_ > f_Combo_*& *f_W_*). For lines evolved in CdCu, CoCu, and CoMn, fitness was significantly higher in the combination environment than in either single metal (*f_Combo_ > f_S_*& *f_W_*), indicating that rescue in the combined metal environment provided less benefit for growth in the individual metals. Finally, lines evolved in NiZn exhibited equivalently high fitness in the combination environment and the stronger interactor, Ni, and lower fitness in the weaker interactor, Zn (*f_Combo_*& *f_S_ > f_W_*).

**Table 1:**
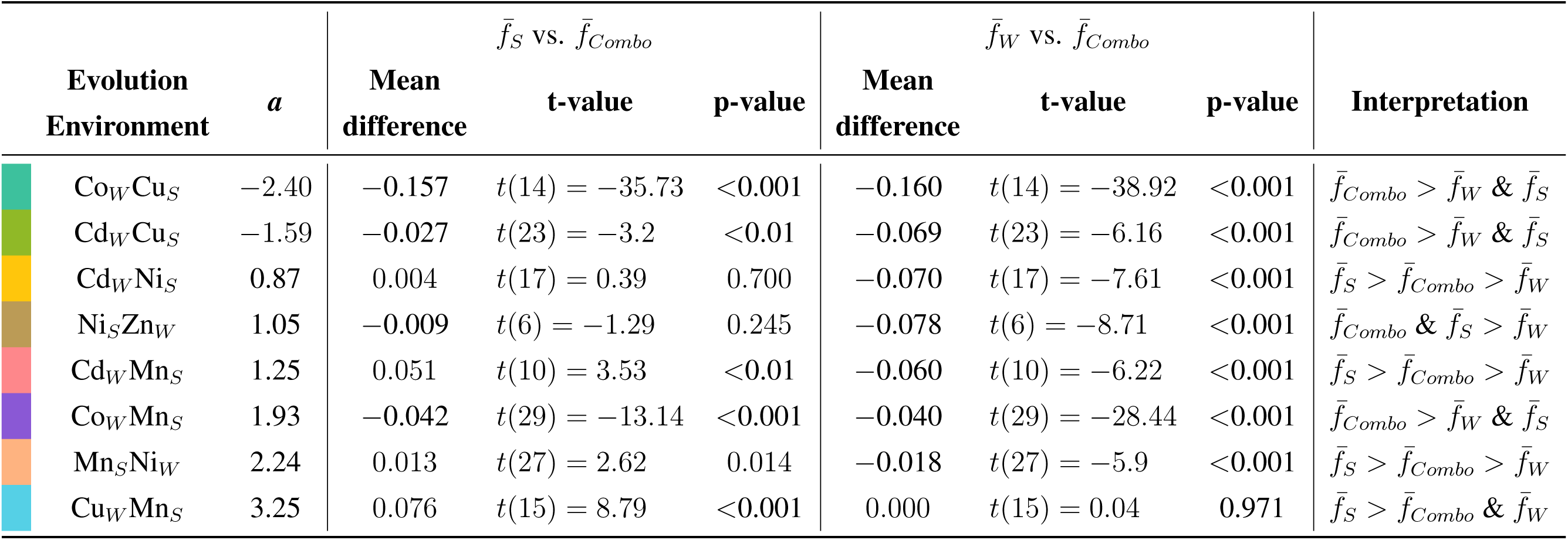
Paired t-tests comparing relative fitness in single metals vs. combination environments. For “Mean difference”, the reference is *f̄Combo*. We consider p values below *α* = 0.025 to be significant, considering the two tests conducted for that metal combination.

We were unable to test the effect of the respiratory phenotype on fitness patterns in many of the metal combinations due to a lack of variation (e.g., all 15 CoCu lines were petite, 6 of 7 NiZn lines were grande; Figure 1B). We ran two-way ANOVAs on fitnesses of grandes and petites for combinations with more than one evolved line in the minority respiratory phenotype (i.e., CdCu, CdNi, CoMn, and MnNi; Table 2, left). While the separate effects of test environment and respiratory phenotype were both significant for fitness of lines evolved in CdNi and MnNi, their interaction was not, meaning that the fitness difference between grandes and petites was similar across the four test environments. The interaction was significant for fitness of lines evolved in CdCu and CoMn. Tukey post-hoc tests (Table 2, right) revealed that the CoMn petites grew worse than the grandes in YPAD and the stronger interactor, Mn, with similar fitness in the metal combination and Co. The CdCu petites also had lower fitness in YPAD, however, the petites grew significantly better than the grandes in the metal combination and in the weaker interactor, Cd. These results suggest that petites may have contributed to evolutionary rescue, but only in some metals, a point that we return to in the Discussion.

**Table 2:**
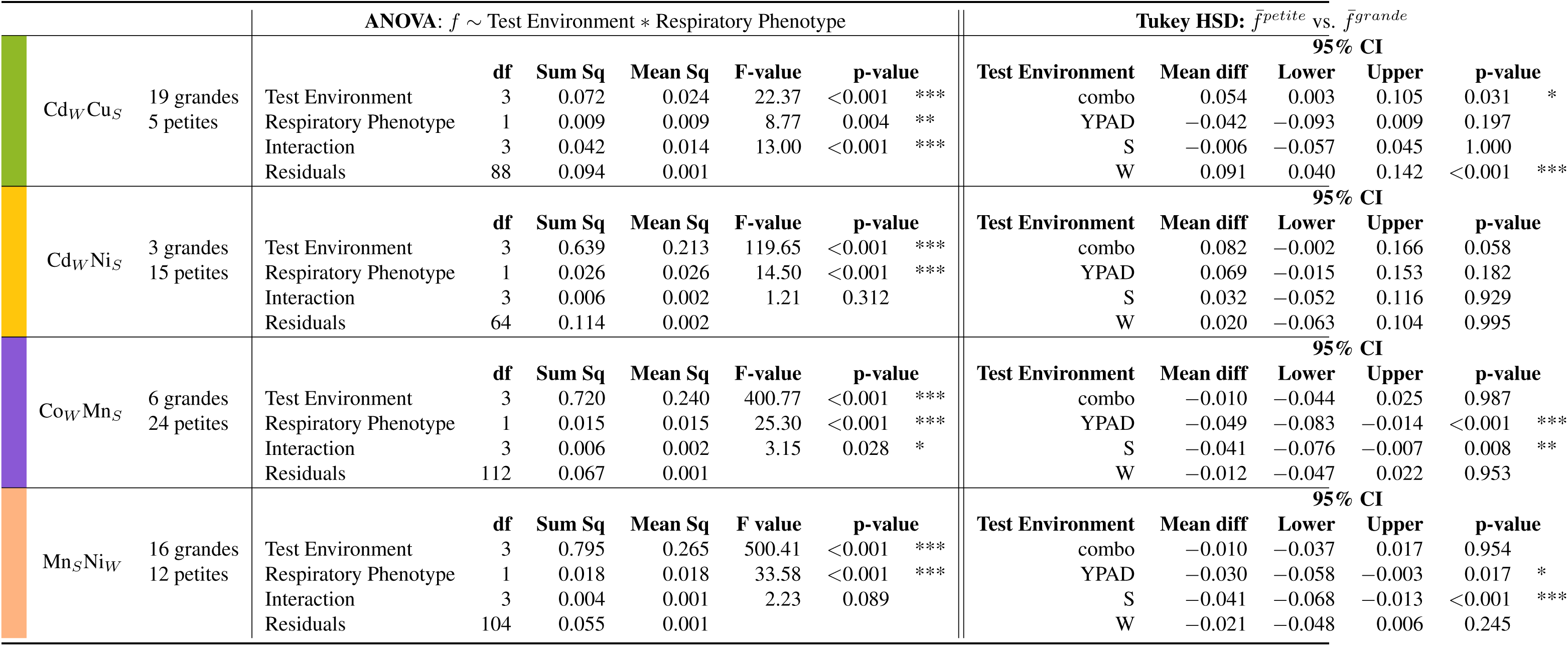
Comparing relative fitness (*f*) of petite and grande lines in each test environment. Comparisons are only made for combinations with more than 1 line in the minority respiratory phenotype. (left) Two way ANOVAs analyzing the effect of test environment and respiratory phenotype on relative fitness of grandes vs. petites. The model is given in the left header. (right) Tukey post-hoc test showing the comparisons between average relative fitness of petites (*f̄^petite^*) and average relative fitness of grandes (*f̄^grande^*) in each test environment.

| | | ANOVA: $f \sim$ Test Environment * Respiratory Phenotype | | | | | | Tukey HSD: $\bar{f}^{petite}$ vs. $\bar{f}^{grande}$ | | | | |
| --- | --- | --- | --- | --- | --- | --- | --- | --- | --- | --- | --- | --- |
|  |  |  | df | Sum Sq | Mean Sq | F-value | p-value | Test Environment | Mean diff | 95% CI |  | p-value |
|  | Cd<sub>W</sub>Cu<sub>S</sub> 19 grandes<br>5 petites | Test Environment | 3 | 0.072 | 0.024 | 22.37 | <0.001 *** | combo | 0.054 | 0.003 | 0.105 | 0.031 |
|  |  | Respiratory Phenotype | 1 | 0.009 | 0.009 | 8.77 | 0.004 ** | YPAD | -0.042 | -0.093 | 0.009 | 0.197 |
|  |  | Interaction | 3 | 0.042 | 0.014 | 13.00 | <0.001 *** | S | -0.006 | -0.057 | 0.045 | 1.000 |
|  |  | Residuals | 88 | 0.094 | 0.001 |  |  | W | 0.091 | 0.040 | 0.142 | <0.001 |
|  | Cd<sub>W</sub>Ni<sub>S</sub> 3 grandes<br>15 petites |  | df | Sum Sq | Mean Sq | F-value | p-value | Test Environment | Mean diff | 95% CI |  | p-value |
|  |  | Test Environment | 3 | 0.639 | 0.213 | 119.65 | <0.001 *** | combo | 0.082 | -0.002 | 0.166 | 0.058 |
|  |  | Respiratory Phenotype | 1 | 0.026 | 0.026 | 14.50 | <0.001 *** | YPAD | 0.069 | -0.015 | 0.153 | 0.182 |
|  |  | Interaction | 3 | 0.006 | 0.002 | 1.21 | 0.312 | S | 0.032 | -0.052 | 0.116 | 0.929 |
|  |  | Residuals | 64 | 0.114 | 0.002 |  |  | W | 0.020 | -0.063 | 0.104 | 0.995 |
|  | Co<sub>W</sub>Mn<sub>S</sub> 6 grandes<br>24 petites |  | df | Sum Sq | Mean Sq | F-value | p-value | Test Environment | Mean diff | 95% CI |  | p-value |
|  |  | Test Environment | 3 | 0.720 | 0.240 | 400.77 | <0.001 *** | combo | -0.010 | -0.044 | 0.025 | 0.987 |
|  |  | Respiratory Phenotype | 1 | 0.015 | 0.015 | 25.30 | <0.001 *** | YPAD | -0.049 | -0.083 | -0.014 | <0.001 |
|  |  | Interaction | 3 | 0.006 | 0.002 | 3.15 | 0.028 * | S | -0.041 | -0.076 | -0.007 | 0.008 |
|  |  | Residuals | 112 | 0.067 | 0.001 |  |  | W | -0.012 | -0.047 | 0.022 | 0.953 |
|  | Mn<sub>S</sub>Ni<sub>W</sub> 16 grandes<br>12 petites |  | df | Sum Sq | Mean Sq | F value | p-value | Test Environment | Mean diff | 95% CI |  | p-value |
|  |  | Test Environment | 3 | 0.795 | 0.265 | 500.41 | <0.001 *** | combo | -0.010 | -0.037 | 0.017 | 0.954 |
|  |  | Respiratory Phenotype | 1 | 0.018 | 0.018 | 33.58 | <0.001 *** | YPAD | -0.030 | -0.058 | -0.003 | 0.017 |
|  |  | Interaction | 3 | 0.004 | 0.001 | 2.23 | 0.089 | S | -0.041 | -0.068 | -0.013 | <0.001 |
|  |  | Residuals | 104 | 0.055 | 0.001 |  |  | W | -0.021 | -0.048 | 0.006 | 0.245 |

### Correlation between fitnesses in different test environments

We next investigated if lines exhibited different fitness trade-offs in the different test environments when evolved in a specific metal combination. Different trade-offs may point to underlying mutations affecting different structures or pathways. Specifically, we looked for significant correlations between relative fitnesses in the four different test environments (metal combination, YPAD, stronger interactor, and weaker interactor). Figure 5 displays the significance of Pearson correlation coefficients between fitness in different test environments for lines evolved in each combination (evolution environment given below each grid, with stronger and weaker interactors given as subscripts to each individual metal). Fitness measurements of evolved lines (red for petite and blue for grande) are overlain on heatmaps showing the significance of the correlations, with lighter yellow representing lower significant *p*-values and white representing non-significance (*p >* 0.05).

**Figure 5:**
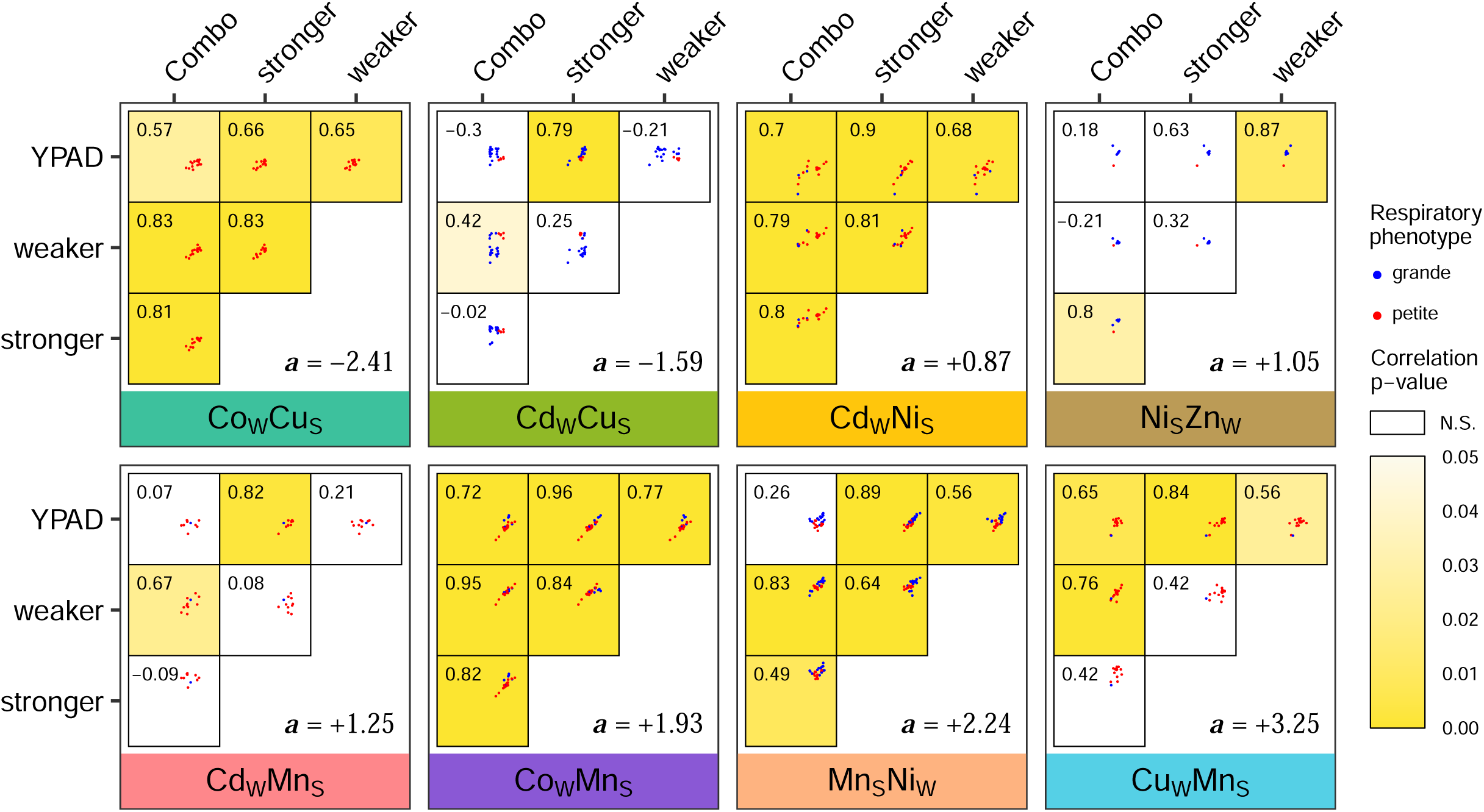
Correlations between relative fitnesses in all test environments for each metal combination. Pairwise grids for each metal combination (identified at bottom of each grid) showing correlations between relative fitnesses in all test environments. Each cell within a grid shows the following: a scatterplot with relative fitnesses in two environments, with red dots representing petites and blue dots representing grandes (x-axis is environment at top, y-axis is environment at left), with identical axis scales in all cells and grids; Pearson’s correlation coefficient is given in the top left; and a color representing significance of the correlation, with non-significant correlations in white and significant correlations in shades of yellow.

Overall, correlations in the growth rates between two environments were either significantly positive among lines (33 comparisons) or non-significant (15 comparisons). The absence of significantly negative correlations means that lines with most-improved fitness in the metal combination did not trade off this fitness gain with much lower fitness in YPAD or the single metals. Rather, lines rescued in a particular metal combination largely exhibited similar fitness effects across all environments. For example, lines evolved in CdNi (N = 20), CoCu (N = 17), and CoMn (N = 30) have significantly positive correlations in growth among all tested environments. That is, some rescue lines were more fit across all environments tested than other lines. In other cases, lines were similar in fitness in different environments and so did not reveal significant relationships (clear boxes in Figure 5).

## Discussion

Our findings demonstrate a large diversity in the fitness consequences of Saccharomyces cerevisiae populations that have undergone evolutionary rescue in heavy metal combinations. We found that rescue was more common in metal combinations that are less toxic when combined than expected from the component metals (”antagonistic” combinations), and in these environments, adaptation increased resistance to individual metals, as well as the metal combination. We also observed rescue in two metal combinations that are more toxic when combined than expected (”synergistic”), but in these cases evolved lines tended to have more specialized adaptations to the toxic effects of the metal combination and fitness often declined in the component metals. By contrast, we never observed evolutionary rescue in metal combinations where toxicity of the two metals combined additively.

We initially hypothesized that strongly interacting metal combinations would hinder rescue by limiting mutations to the subset able to resolve the particular harms caused by the metal interactions. Nevertheless, we observed rescue in both antagonistic and synergistic metal combinations.

In antagonistic stressor combinations, the concentration-response surface of the ancestor is flatter (see black curves in Figure 2A and Kahn and Otto (2025)), allowing a broader range of mutations to increase fitness. Here, different stressors may act on (or be regulated by) distinct yet interconnected cellular pathways, such that the response to either stressor is favourable, potentially reducing conflicts in resource allocation or regulatory responses and permitting mutations that confer resistance without an overwhelming fitness cost Goussen et al. (2020).

In synergistic combinations, the concentration-response surface of the ancestor steeply declines as the two metals combine, indicating an emergent harm caused by their interaction (see black curves in Figure 2A). While this may restrict opportunities for rescue to that subset of single-step mutations able to tolerate this harm, such resistance mutations may have a larger selective advantage Hegreness et al. (2008), resolving the emergent harm at concentrations where the single metals are not that toxic.

Opposite to our hypothesis, we observed no rescue events in any of the three metal combinations that were consistent with concentration-addition (CdCo, CdZn, or CoZn with *a* not significantly different from 0). These metals act like one another in that their concentration-response surfaces are shaped as though the two metals were the same. The metals are clearly not equivalent from a cellular perspective in these cases, however, as we would then expect rescue as often as observed in the single metals Kahn and Otto (2025). Rather, the metal combinations obeying concentration addition may have a mix of emergent harms and single metal stresses that make it challenging to rescue. If different mutations, specific to each metal and their combination, are required for rescue, the chances of them arising in the same cell may be too low to occur before extinction, or they may simply be incompatible or inviable when combined (e.g., sign epistasis Zhang et al. (2024), reciprocal sign epistasis Chiotti et al. (2014)), requiring further compensatory mutations to reconcile their conflicting effects.

However, it is difficult to disentangle whether the observed differences in rescue frequency across combinations with different interaction types are general properties of such interactions or are instead artifacts of the specific metal concentrations used. Even small deviations in metal concentrations could shift populations across thresholds between acclimation, evolutionary rescue, and inevitable extinction. This uncertainty highlights a common challenge in experimental studies of evolutionary rescue, where a fine balance of environmental stress is required to enable rescue but not growth Carlson et al. (2014). Future work incorporating finer gradations of stressor intensity, larger population sizes, and longer selection periods could help determine whether some combinations truly constrain evolutionary rescue, or whether rescue in those combinations is possible but was simply not captured in these experiments.

Without yet having whole-genome sequences, we can only speculate on the underlying genetic and cellular mechanisms for the observed fitness patterns at this stage; however, these data will be incorporated and analyzed in detail in the near future.

Analyzing growth rates in single metals and their combination, as well as YPAD, we found that adaptations for antagonistic combinations tended to increase fitness in each individual metal, whereas evolution in synergistic combinations led to more specialized adaptations, with fitness benefits largely confined to the combination environment and fitness costs in the single metals (Figures 2, 3). This pattern suggests that antagonistic interactions may select for mutations with more general benefits to growth (e.g., decreasing inhibition of the cell cycle), while synergistic interactions may favour mutations tailored to emergent properties arising in the combination, with fitness benefits largely confined to the combination environment.

We next speculate about the cellular mechanisms underlying these patterns by considering the physical and chemical properties of the most strongly interactive metals, manganese (antagonistic driver) and copper (synergistic driver).

Cells evolving in combinations with manganese, which can form antioxidant complexes with small molecules Reddi and Culotta (2013), likely experience less oxidative burden, allowing selection to act more directly on the non-oxidative components of metal stress. When tested in the weaker interactor, these cells benefit somewhat from resistance alleles for the non-oxidative components of stress, increasing their fitness relative to the ancestor in this environment (except for CdMn lines, Figure 3D). However, significantly lower fitness in the weaker interactor compared to in the combination, as is the case for lines evolved in CdMn, CoMn, and MnNi (Table 1), implies that, without manganese, these cells cannot withstand the full brunt of the toxic burden imposed by the weaker interactor on its own. Additionally, significantly higher fitness in manganese compared to in the combination, as is the case for lines evolved in CdMn, CuMn, and MnNi (Table 1), demonstrates that these lines are particularly well-adapted to manganese, suggesting that mutations favourable in manganese remain favourable in the combination.

Cells evolved in combinations with copper, which permeabilizes the plasma membrane Gaetke (2003) and directly produces ROS via Fenton-like reactions Macomber and Imlay (2009), likely experience increased uptake of both metals and increased oxidative stress. For lines evolved in the two environments including copper (i.e., CdCu and CoCu), relative fitnesses in the single metals were significantly lower than fitness in the combination (Table 1), indicating that the adaptations arising in these environments are not targeted for the effects of the individual metals, but for some emergent property that arises due to the combination. This echoes patterns seen in drug resistance evolution, where exposure to multi-drug environments with synergistic toxicity can select for highly specific resistance mechanisms that do not necessarily confer cross-resistance to individual drugs Hegreness et al. (2008); Chevereau and Bollenbach (2015).

Mitochondria, which intrinsically produce ROS during respiration Carmona-Gutierrez et al. (2010), contribute to the overall oxidative burden in the cell, which may be exacerbated by the presence of metals, but partially alleviated by loss of the organelle. Given that both cadmium and copper induce high levels of oxidative stress compared to the other metals used in this experiment Valko et al. (2005), loss of mitochondria and the accompanying decrease in ROS could be particularly advantageous. Indeed, the petite lines from CdCu had significantly higher fitness compared to the grandes in both the combination environment and in cadmium on its own (Table 2), pointing to an advantage of the petite phenotype in these metals. Cadmium has been shown to directly bind and inhibit function of essential components of the electron transport chain Miccadei and Floridi (1993); Dorta et al. (2003), as well as increase permeability of the inner mitochondrial membrane, dissipating the membrane potential required for electron transfer Kravens’ka et al. (2008). Our growth curves from yeast grown in cadmium also show evidence of mitochondrial dysfunction. The maximum OD in cadmium is around 0.8, compared to ODs over 1.0 in all other metals tested Kahn and Otto (2025). Function of the mitochondria is fundamental to yeast’s ability to switch metabolism glycolysis to respiration. Once once the fermentable sugars are consumed, the yeast must switch to aerobic utilization of ethanol, which is what accounts for reduced (but still positive) growth rates at later phases of the culture Galdieri et al. (2010). While the mitotoxic effects of cadmium suggest that petites may be favoured in cadmium environments, many of the rescue lines in CdCu were not petites (Figure 1B). Data obtained from whole-genome sequencing of these lines will clarify the role of mitochondrial loss versus other genetic mechanisms in adaptation.

Throughout history, organisms have been subject to multiple selection pressures simultaneously, but human-driven activities are exposing the natural world to an increasing array of stressor combinations, the components of which can interact to increase or decrease the sum of their harmful effects. These interactions can change the trajectory of evolutionary responses, as we have documented. Understanding how stressor interactions shape evolution will become increasingly important as environmental conditions continue to intensify. Our results highlight that it is not only the presence or intensity of individual stressors that determines evolutionary outcomes, but also the way these stressors interact, whether they amplify, buffer, or create new, emergent selection pressures. Our findings suggest that evolutionary rescue may be less likely in metal combinations exhibiting concentration addition compared to interactive combinations, regardless of how the interaction affects stressor intensity. Surprisingly, while synergistic metal interactions may be more harmful in terms of mortality, they may not be worse in terms of adaptive potential.

## Acknowledgements

We are grateful J Grant and K Bakhtari Mousakhan for their help in data collection, to M Whitlock, R Redfield, S Baldwin, and A Leeks for their feedback and comments on the manuscript. Funding was provided by the Natural Sciences and Engineering Research Council of Canada (RGPIN-2022*−*03726 to SPO) and a four-year fellowship from the University of British Columbia (to PCK).

## Author Contributions

PCK conceived the concept and experimental design, collected and analyzed the data, produced the figures, and wrote the manuscript. SPO INSERT HERE

