## Supplemental Material for "Evolutionary rescue in toxic combinations of metal is dependent on metal interactions"

Journal: *bioRxiv*

Manuscript type: *Preprint*

[2]Present Address: *Department of Microbiology, University of Manitoba, Canada*

#### Supplementary Methods

Determining concentrations for evolutionary rescue in metal combinations Metal concentrations for evolutionary rescue in binary metal combinations were chosen with two goals: (i) equal toxicity from each individual metal and (ii) a combined toxicity just past the threshold allowing growth, requiring the emergence of beneficial alleles to rescue the line. Because interactions between metals directly affects the toxicity of the environment (Chapter 3), evolutionary rescue required higher concentrations in antagonistic combinations and lower concentrations in synergistic combinations. We sought to determine how much higher or lower the concentrations needed to be for each combination.

To infer these concentrations, checkerboard assays, which are traditionally used to test the effects of drug interactions on microbial growth were performed (method described in detail by Bellio et al., 2021). Briefly, for a combination of metals A and B, a gradient of eight concentrations of metal A was arrayed along the rows of a BioTek plate, and a gradient of eight concentrations of metal B was arrayed along the columns, creating a matrix of concentration pairs (Figure S4.1A); the gradients of each individual metal were also tested in isolation (x and y axes of Figure S4.1A; numerical values of OD and concentration are shown in Figure S4.1B). Plates were sealed with clear adhesive film and shaken at 30 ◦C for 48 hours, after which the film was removed, and OD was measured in a BioTek spectrophotometer. 48 hours was chosen as the measurement period to simulate the first days of the evolutionary rescue experiment, during which wells should not display growth.

Nine checkerboard assays were conducted, ensuring representation of all six metals and a range of interaction effects. As we had not yet determined a values when these assays were conducted, qualitative categories of growth rates observed in Figure 3.1A were used to group all 15 possible metal combinations into four groups: (1) ”growth in combo $\geq$ growth in singles” (combos MnZn and CuMn with a checkerboard assay of MnZn); (2) ”growth in combo ¡ growth in singles” (combos CdMn, CoMn, and MnNi with checkerboard assays of CdMn and CoMn);

(3)”growth in combo $\gg$ growth in singles” (combos CdCo, CdCu, CdNi, CdZn, CoNi, CoZn, and NiZn with checkerboard assays of CdZn, CoNi, CoZn, and NiZn). (4) ”growth in combo $\approx$ 0”, that is, undetectable by spectrophotometry (combos CoCu, CuNi, and CuZn with a checkerboard assay of CuNi).

Through analysis of the checkerboard assay results (Figure S4.1), we determined concentrations of single metals that produced equivalent OD measurements after 48 hours and, when combined, no observable growth. For combinations in group 1, concentrations of each single metal ([A] and [B]) producing a OD48hr of 0.0-0.1 prevent observable growth when combined. For combinations in group 2, [A] and [B] producing an OD48hr $\sim$0.3 prevent observable growth when combined. For combinations in group 3, [A] and [B] producing an OD48hr $\sim$0.6-0.7 prevent observable growth when combined. For combinations in group 4, [A] and [B] producing an OD48hr $\sim$0.8-0.9 or higher prevent observable growth when combined. Using those target ranges, the concentrations of each individual metal were chosen for evolutionary rescue in each combination and are shown in Table S1. These targets were presumed to be effective for combinations that were not assayed via checkerboard because similar maximum growth rates were observed among combinations within the same qualitative grouping. We recognize that variation within qualitative groupings may have led to suboptimal concentration choices for some combinations, preventing evolutionary rescue in those environments.

### Supplementary Tables

Table 1: Metal concentrations: (A) Concentrations (mM) of individual metals used in the evolutionary rescue experiments and number of wells with growth by the end of the 14-day period. Concentrations were chosen based on checkerboard assay results, aiming for equivalent $OD_{48hr}$ in the individual metals and no observable growth after 48 hours in the combination. (B) Concentrations of individual metals used in the fitness assays. Concentrations were chosen based on v-graph assays and concentration-response curves (**?**, aiming for equivalent maximum growth rates in the individual metals and a max. growth rate of 0.05 in the combination.

**(A) Evolution experiments**

| Combo | Qualitative category | Single metal target OD | Metal | Trial 1 conc (mM) | Trial 1 Wells with growth | Trial 2 conc (mM) | Trial 2 Wells with growth |
|---|---|---|---|---|---|---|---|
| CdCo | 3 | 0.6-0.7 | Cd | 0.0082 | 80 | 0.015 | 80 |
| | | | Co | 1.14 | | 1.14 | |
| CdCu | 3 | 0.6-0.7 | Cd | 0.0082 | **24** | | |
| | | | Cu | 11.3 | | | |
| CdMn | 2 | 0.3-0.4 | Cd | 0.015 | 80 | 0.022 | **11** |
| | | | Mn | 2.95 | | 3.4 | |
| CdNi | 3 | 0.6-0.7 | Cd | 0.015 | **20** | | |
| | | | Ni | 3.0 | | | |
| CdZn | 3 | 0.6-0.7 | Cd | 0.0082 | 80 | 0.015 | 0 |
| | | | Zn | 4.2 | | 5.15 | |
| CoCu | 4 | 0.9-1.0 | Co | 0.6 | 0 | 0.51 | **17** |
| | | | Cu | 10.0 | | 8.5 | |
| CoMn | 2 | 0.3-0.4 | Co | 1.5 | **30** | | |
| | | | Mn | 2.95 | | | |
| CoNi | 3 | 0.6-0.7 | Co | 1.14 | 80 | 1.5 | 0 |
| | | | Ni | 2.0 | | 3.0 | |
| CoZn | 3 | 0.6-0.7 | Co | 1.14 | 0 | 0.96 | 0 |
| | | | Zn | 4.2 | | 3.57 | |
| CuMn | 1 | 0.0-0.1 | Cu | 12.0 | **16** | | |
| | | | Mn | 2.95 | | | |
| CuNi | 4 | 0.9-1.0 | Cu | 10.0 | 0 | 8.5 | 0 |
| | | | Ni | 1.4 | | 1.19 | |
| CuZn | 4 | 0.9-1.0 | Cu | 10.0 | 0 | 8.5 | 0 |
| | | | Zn | 2.25 | | 1.91 | |
| MnNi | 2 | 0.3-0.4 | Mn | 2.95 | **28** | | |
| | | | Ni | 3.0 | | | |
| MnZn | 1 | 0.0-0.1 | Mn | 3.85 | 0 | 3.27 | 0 |
| | | | Zn | 10.0 | | 8.5 | |
| NiZn | 3 | 0.6-0.7 | Ni | 2.0 | **7** | | |
| | | | Zn | 4.2 | | | |

**(B) Fitness assays**

| combo | metal | conc (mM) |
|---|---|---|
| CoCu | Co | 0.35 |
| | Cu | 6.0 |
| CdCu | Cd | 0.015 |
| | Cu | 11.25 |
| CdNi | Cd | 0.037 |
| | Ni | 2.25 |
| NiZn | Ni | 1.86 |
| | Zn | 2.0 |
| CdMn | Cd | 0.034 |
| | Mn | 0.82 |
| CoMn | Co | 0.85 |
| | Mn | 0.80 |
| MnNi | Mn | 0.94 |
| | Ni | 2.25 |
| CuMn | Cu | 13.5 |
| | Mn | 0.8 |

Table 2: Cell density: Estimating cell density of OLY077 overnight culture. (A) Estimation of cells/mL in overnight culture (incubated in liquid YPAD shaking for 24 hour at 30 °C) from hemocytometer counts of 200x dilution. Equation for calculation: $Average\ cells\ per\ square \times Dilution\ factor \times 10^4$. 8 squares were counted per trial. (B) Dilution Series: Cell densities estimated from dilution series with OD measurements. A linear fit to this data was performed in R using the texttlm() function. (C) Estimating number of cells in founding populations. Equation of linear fit to data in part B: $cells/mL = 198,529,638 \times OD - 21,970,423$.

**(A) Hemocytometer summary**

| | Trial 1 | Trial 2 | Trial 3 | Average |
|---|---|---|---|---|
| Average cells per square: | 134 | 113.375 | 149.875 | 132 |
| Cells/mL undiluted culture: | 268000000 | 226750000 | 299750000 | 264833333 |

**(B) Dilution series**

| Dilution Factor | OD | Est. cells/mL |
|---|---|---|
| 1 | 1.140 | 264833333 |
| 2 | 0.760 | 132416667 |
| 4 | 0.469 | 66208333 |
| 6 | 0.344 | 44138889 |
| 8 | 0.286 | 33104167 |
| 10 | 0.242 | 26483333 |
| 16 | 0.186 | 16552083 |
| 20 | 0.166 | 13241667 |
| 100 | 0.086 | 2648333 |
| 200 | 0.112 | 1324167 |

**(C) Estimating cell counts of founding populations**

| | Normalized culture | | | | | |
|---|---|---|---|---|---|---|
| | OD | Estimated cells/mL (uL) | 1:10 cells/mL | cells/$\mu$L | Inoculum volume ($\mu$L) | cells/inoculum |
| Evolution experiment | 0.2 | 17735505 | 1773551 | 1774 | 10 | 17740 |
| Fitness assay | 0.7 | 117000324 | 11700032 | 11700 | 9 | 105300 |

Table 3: Evolved lines: Information on evolved lines including plate-well location of rescue, date first observed (putative growth if observed before sample date), date sampled (”original sample”), and date of confirmed growth (OD ¿ 0.5 in 2 mL metal media); respiratory phenotype; and relative fitnesses in each test environment (average of three technical replicates). Lines excluded because of failure to grow to OD = 0.5 in four days after sampling are highlighted in red.

| trial | combo | line | well | date observed | date sampled | date confirmed | petite_test | Relative Fitness ($r_{Evo} - r_{Anc}$ f_Combo | f_YPAD | f_S | f_W |
|---|---|---|---|---|---|---|---|---|---|---|---|
| Trial 1 13-Dec | CdCu | 01 | H10 | 14-Dec | 14-Dec | 15-Dec | grande | -0.010 | -0.023 | 0.008 | -0.097 |
| | | 02 | A9 | 15-Dec | 15-Dec | 16-Dec | grande | -0.014 | -0.003 | 0.008 | -0.107 |
| | | 03 | A11 | 14-Dec | 15-Dec | 16-Dec | grande | -0.018 | -0.019 | 0.013 | -0.061 |
| | | 04 | C7 | 15-Dec | 15-Dec | 16-Dec | grande | 0.032 | -0.016 | 0.006 | -0.067 |
| | | 05 | D11 | 14-Dec | 15-Dec | 16-Dec | grande | 0.024 | -0.014 | -0.010 | -0.110 |
| | | 06 | B11 | 16-Dec | 16-Dec | 17-Dec | grande | -0.007 | -0.046 | 0.006 | 0.026 |
| | | 07 | C6 | 15-Dec | 16-Dec | 17-Dec | grande | 0.027 | -0.049 | -0.013 | -0.072 |
| | | 08 | D9 | 15-Dec | 16-Dec | 17-Dec | grande | -0.009 | -0.021 | 0.005 | -0.083 |
| | | 09 | G10 | 15-Dec | 16-Dec | 17-Dec | grande | -0.015 | -0.064 | -0.019 | -0.112 |
| | | 10 | H4 | 15-Dec | 16-Dec | 17-Dec | grande | 0.033 | -0.067 | -0.030 | -0.087 |
| | | 11 | H11 | 15-Dec | 16-Dec | 17-Dec | grande | -0.018 | -0.047 | 0.004 | 0.014 |
| | | 12 | A3 | 17-Dec | 17-Dec | 18-Dec | grande | -0.013 | -0.016 | 0.006 | -0.097 |
| | | 13 | A8 | 17-Dec | 17-Dec | 18-Dec | grande | -0.004 | -0.018 | 0.001 | -0.100 |
| | | 14 | B9 | 16-Dec | 17-Dec | 18-Dec | grande | 0.021 | -0.013 | 0.006 | 0.027 |
| | | 15 | E8 | 16-Dec | 17-Dec | 18-Dec | grande | -0.004 | -0.092 | -0.092 | -0.080 |
| | | 16 | F5 | 16-Dec | 17-Dec | 18-Dec | petite | 0.070 | -0.069 | -0.006 | -0.005 |
| | | 17 | B3 | 17-Dec | 18-Dec | 19-Dec | grande | -0.019 | -0.034 | -0.008 | -0.006 |
| | | 18 | D3 | 18-Dec | 18-Dec | 19-Dec | petite | 0.052 | -0.078 | -0.021 | 0.016 |
| | | 19 | B5 | 18-Dec | 19-Dec | 21-Dec | grande | -0.016 | -0.118 | -0.102 | -0.163 |
| | | 20 | F9 | 18-Dec | 19-Dec | 20-Dec | grande | 0.036 | -0.013 | -0.007 | -0.105 |
| | | 21 | C2 | 20-Dec | 20-Dec | 21-Dec | petite | 0.040 | -0.088 | -0.015 | 0.026 |
| | | 22 | D4 | 20-Dec | 20-Dec | 21-Dec | petite | 0.053 | -0.083 | -0.024 | 0.021 |
| | | 23 | G5 | 22-Dec | 22-Dec | 23-Dec | petite | 0.070 | -0.077 | -0.022 | 0.029 |
| | | 24 | G9 | 24-Dec | 24-Dec | 25-Dec | grande | 0.029 | -0.032 | -0.007 | -0.122 |
| | | 01 | F5 | 17-Dec | 18-Dec | 19-Dec | petite | 0.106 | -0.137 | 0.080 | 0.011 |
| | | 02 | G10 | 18-Dec | 18-Dec | 19-Dec | petite | 0.162 | -0.099 | 0.129 | 0.062 |
| | | 03 | A9 | 18-Dec | 19-Dec | 21-Dec | petite | 0.119 | -0.142 | 0.088 | 0.017 |
| | | 04 | B3 | 19-Dec | 19-Dec | 21-Dec | petite | 0.105 | -0.148 | 0.080 | 0.013 |
| | | 05 | B10 | 19-Dec | 19-Dec | 20-Dec | petite | 0.113 | -0.141 | 0.089 | 0.016 |
| | | 06 | C4 | 18-Dec | 19-Dec | 21-Dec | petite | -0.019 | -0.243 | 0.010 | -0.052 |
| | | 07 | F11 | 19-Dec | 19-Dec | 21-Dec | grande | -0.020 | -0.307 | 0.017 | -0.043 |
| | | 08 | G7 | 18-Dec | 19-Dec | 21-Dec | petite | 0.099 | -0.136 | 0.085 | 0.005 |
| | | 09 | H4 | 18-Dec | 19-Dec | 21-Dec | petite | -0.012 | -0.202 | 0.035 | -0.046 |
| | | 10 | A2 | 18-Dec | 20-Dec | 21-Dec | grande | 0.039 | -0.159 | 0.065 | 0.046 |

| | | | | | | | | | | | |
|---|---|---|---|---|---|---|---|---|---|---|---|
| Trial 1 13-Dec | CdNi | 11 | E2 | 18-Dec | 20-Dec | 21-Dec | petite | 0.114 | -0.181 | 0.060 | 0.009 |
| | | 12 | E7 | 19-Dec | 20-Dec | 22-Dec | petite | 0.101 | -0.148 | 0.082 | 0.027 |
| | | 13 | D8 | 19-Dec | 21-Dec | 23-Dec | petite | 0.014 | -0.168 | 0.066 | -0.033 |
| | | 14 | G8 | 19-Dec | 21-Dec | 23-Dec | petite | 0.040 | -0.128 | 0.056 | -0.045 |
| | | 15 | H9 | 20-Dec | 21-Dec | 23-Dec | grande | -0.016 | -0.185 | 0.057 | -0.055 |
| | | 16 | A4 | 20-Dec | 23-Dec | 24-Dec | petite | 0.147 | -0.112 | 0.105 | 0.026 |
| | | 17 | G2 | 22-Dec | 23-Dec | 24-Dec | petite | 0.106 | -0.143 | 0.091 | -0.001 |
| | | 18 | C7 | 22-Dec | 25-Dec | 26-Dec | petite | 0.048 | -0.093 | 0.118 | 0.037 |
| | | 19 | H2 | 23-Dec | 25-Dec | 30-Dec | NA | NA | NA | NA | NA |
| | | 20 | H3 | 23-Dec | 25-Dec | 30-Dec | NA | NA | NA | NA | NA |
| Trial 1 13-Dec | CoMn | 01 | B4 | 15-Dec | 18-Dec | 19-Dec | petite | 0.097 | -0.118 | 0.039 | 0.059 |
| | | 02 | D8 | 17-Dec | 18-Dec | 20-Dec | petite | 0.078 | -0.121 | 0.035 | 0.032 |
| | | 03 | F5 | 19-Dec | 20-Dec | 21-Dec | grande | 0.104 | -0.052 | 0.101 | 0.070 |
| | | 04 | G7 | 18-Dec | 20-Dec | 21-Dec | petite | 0.093 | -0.117 | 0.048 | 0.046 |
| | | 05 | A2 | 20-Dec | 21-Dec | 22-Dec | petite | 0.099 | -0.111 | 0.054 | 0.061 |
| | | 06 | B5 | 20-Dec | 21-Dec | 22-Dec | petite | 0.094 | -0.114 | 0.047 | 0.046 |
| | | 07 | B6 | 20-Dec | 21-Dec | 23-Dec | petite | 0.097 | -0.120 | 0.035 | 0.048 |
| | | 08 | C7 | 20-Dec | 21-Dec | 23-Dec | petite | 0.100 | -0.085 | 0.073 | 0.079 |
| | | 09 | D3 | 20-Dec | 21-Dec | 23-Dec | petite | 0.087 | -0.118 | 0.031 | 0.052 |
| | | 10 | D6 | 20-Dec | 21-Dec | 22-Dec | petite | 0.088 | -0.129 | 0.043 | 0.056 |
| | | 11 | E7 | 20-Dec | 21-Dec | 23-Dec | petite | 0.021 | -0.200 | -0.034 | -0.030 |
| | | 12 | F10 | 20-Dec | 21-Dec | 22-Dec | petite | 0.096 | -0.107 | 0.051 | 0.058 |
| | | 13 | H5 | 21-Dec | 21-Dec | 22-Dec | grande | 0.126 | -0.098 | 0.075 | 0.076 |
| | | 14 | A7 | 21-Dec | 22-Dec | 24-Dec | grande | 0.101 | -0.055 | 0.094 | 0.064 |
| | | 15 | A11 | 22-Dec | 22-Dec | 24-Dec | petite | 0.092 | -0.122 | 0.041 | 0.051 |
| | | 16 | B2 | 21-Dec | 22-Dec | 24-Dec | petite | 0.099 | -0.108 | 0.037 | 0.045 |
| | | 17 | B7 | 22-Dec | 22-Dec | 24-Dec | petite | 0.098 | -0.119 | 0.053 | 0.056 |
| | | 18 | E6 | 22-Dec | 22-Dec | 24-Dec | petite | 0.069 | -0.119 | 0.018 | 0.038 |
| | | 19 | E8 | 20-Dec | 22-Dec | 24-Dec | petite | 0.144 | -0.083 | 0.085 | 0.103 |
| | | 20 | F4 | 21-Dec | 22-Dec | 24-Dec | grande | 0.105 | -0.042 | 0.110 | 0.058 |
| | | 21 | F7 | 21-Dec | 22-Dec | 24-Dec | petite | 0.124 | -0.090 | 0.079 | 0.079 |
| | | 22 | C5 | 21-Dec | 23-Dec | 25-Dec | grande | 0.074 | -0.073 | 0.070 | 0.049 |
| | | 23 | C6 | 21-Dec | 23-Dec | 25-Dec | petite | 0.092 | -0.120 | 0.045 | 0.052 |
| | | 24 | E9 | 22-Dec | 23-Dec | 25-Dec | grande | 0.080 | -0.116 | 0.036 | 0.048 |
| Trial 1 13-Dec | CoMn | 25 | D7 | 22-Dec | 24-Dec | 26-Dec | petite | 0.041 | -0.179 | -0.014 | 0.003 |
| | | 26 | D11 | 23-Dec | 24-Dec | 25-Dec | petite | 0.093 | -0.123 | 0.050 | 0.056 |
| | | 27 | E2 | 23-Dec | 24-Dec | 26-Dec | petite | 0.077 | -0.138 | 0.023 | 0.034 |
| | | 28 | G5 | 23-Dec | 24-Dec | 26-Dec | petite | 0.075 | -0.132 | 0.037 | 0.037 |
| | | 29 | H2 | 21-Dec | 24-Dec | 25-Dec | petite | 0.077 | -0.126 | 0.029 | 0.040 |
| | | 30 | E5 | 25-Dec | 25-Dec | 26-Dec | petite | 0.094 | -0.112 | 0.051 | 0.060 |
| | | 01 | A7 | 17-Dec | 17-Dec | 18-Dec | petite | 0.048 | -0.085 | 0.138 | 0.056 |

| | | | | | | | | | | | |
|---|---|---|---|---|---|---|---|---|---|---|---|
| Trial 1 13-Dec | CuMn | 02 | B11 | 16-Dec | 17-Dec | 18-Dec | petite | 0.047 | -0.087 | 0.156 | 0.055 |
| | | 03 | A10 | 16-Dec | 19-Dec | 20-Dec | petite | 0.073 | -0.077 | 0.139 | 0.077 |
| | | 04 | A3 | 18-Dec | 20-Dec | 22-Dec | petite | 0.008 | -0.169 | 0.058 | -0.002 |
| | | 05 | A6 | 17-Dec | 20-Dec | 22-Dec | petite | 0.043 | -0.064 | 0.141 | 0.046 |
| | | 06 | E5 | 18-Dec | 20-Dec | 22-Dec | petite | 0.024 | -0.070 | 0.146 | 0.023 |
| | | 07 | D11 | 19-Dec | 21-Dec | 26-Dec | NA | NA | NA | NA | NA |
| | | 08 | A11 | 22-Dec | 22-Dec | 23-Dec | petite | 0.046 | -0.079 | 0.147 | 0.060 |
| | | 09 | B10 | 22-Dec | 22-Dec | 23-Dec | petite | 0.036 | -0.085 | 0.141 | 0.064 |
| | | 10 | C5 | 21-Dec | 22-Dec | 23-Dec | petite | 0.052 | -0.091 | 0.060 | 0.036 |
| | | 11 | C10 | 22-Dec | 22-Dec | 23-Dec | petite | 0.042 | -0.077 | 0.137 | 0.050 |
| | | 12 | B7 | 23-Dec | 23-Dec | 25-Dec | grande | 0.011 | -0.172 | 0.037 | 0.009 |
| | | 13 | H10 | 23-Dec | 23-Dec | 24-Dec | petite | 0.049 | -0.107 | 0.091 | 0.045 |
| | | 14 | E4 | 23-Dec | 24-Dec | 26-Dec | petite | 0.059 | -0.061 | 0.156 | 0.058 |
| | | 15 | G2 | 23-Dec | 24-Dec | 25-Dec | petite | 0.032 | -0.109 | 0.117 | 0.033 |
| | | 16 | A4 | 24-Dec | 25-Dec | 26-Dec | petite | 0.046 | -0.092 | 0.135 | -0.013 |
| | | 17 | C7 | 25-Dec | 26-Dec | 30-Dec | petite | 0.079 | -0.091 | 0.106 | 0.097 |
| | | 01 | F5 | 17-Dec | 18-Dec | 19-Dec | grande | 0.138 | -0.034 | 0.159 | 0.135 |
| | | 02 | G11 | 17-Dec | 18-Dec | 19-Dec | grande | 0.120 | -0.103 | 0.101 | 0.093 |
| | | 03 | D6 | 17-Dec | 19-Dec | 21-Dec | grande | 0.123 | -0.049 | 0.147 | 0.117 |
| | | 04 | H11 | 17-Dec | 19-Dec | 21-Dec | grande | 0.142 | -0.042 | 0.159 | 0.110 |
| | | 05 | B7 | 18-Dec | 20-Dec | 21-Dec | grande | 0.128 | -0.057 | 0.149 | 0.123 |
| | | 06 | D10 | 19-Dec | 20-Dec | 21-Dec | grande | 0.129 | -0.050 | 0.151 | 0.123 |
| | | 07 | E11 | 19-Dec | 20-Dec | 21-Dec | petite | 0.140 | -0.098 | 0.129 | 0.110 |
| | | 08 | C9 | 20-Dec | 22-Dec | 24-Dec | grande | 0.114 | -0.070 | 0.147 | 0.117 |
| | | 09 | G6 | 20-Dec | 22-Dec | 24-Dec | petite | 0.082 | -0.092 | 0.110 | 0.092 |
| | | 10 | A4 | 21-Dec | 23-Dec | 26-Dec | grande | 0.106 | -0.061 | 0.136 | 0.104 |
| | | 11 | B4 | 21-Dec | 23-Dec | 25-Dec | petite | 0.094 | -0.103 | 0.077 | 0.086 |
| | | 12 | C3 | 21-Dec | 23-Dec | 24-Dec | petite | 0.102 | -0.115 | 0.092 | 0.073 |
| | | 13 | C4 | 21-Dec | 23-Dec | 24-Dec | petite | 0.108 | -0.141 | 0.078 | 0.067 |
| | | 14 | F10 | 22-Dec | 23-Dec | 26-Dec | grande | 0.056 | -0.065 | 0.131 | 0.028 |
| | | 15 | H2 | 22-Dec | 23-Dec | 24-Dec | petite | 0.106 | -0.112 | 0.104 | 0.080 |
| | | 16 | A7 | 21-Dec | 24-Dec | 25-Dec | petite | 0.132 | -0.106 | 0.124 | 0.091 |
| | | 17 | B8 | 21-Dec | 24-Dec | 26-Dec | petite | 0.088 | -0.122 | 0.099 | 0.079 |
| | | 18 | D8 | 22-Dec | 24-Dec | 26-Dec | petite | 0.130 | -0.108 | 0.122 | 0.083 |
| | | 19 | D11 | 22-Dec | 24-Dec | 26-Dec | grande | 0.059 | -0.077 | 0.126 | 0.058 |
| | | 20 | E7 | 22-Dec | 24-Dec | 26-Dec | grande | 0.136 | -0.051 | 0.156 | 0.125 |
| | | 21 | G10 | 23-Dec | 24-Dec | 26-Dec | grande | 0.143 | -0.027 | 0.181 | 0.143 |
| | | 22 | H4 | 23-Dec | 24-Dec | 26-Dec | grande | 0.071 | -0.081 | 0.112 | 0.056 |
| | | 23 | A8 | 25-Dec | 25-Dec | 26-Dec | grande | 0.141 | -0.083 | 0.133 | 0.116 |
| | | 24 | B2 | 24-Dec | 25-Dec | 26-Dec | petite | 0.090 | -0.075 | 0.124 | 0.084 |
| | | 25 | D9 | 24-Dec | 25-Dec | 26-Dec | grande | 0.151 | -0.071 | 0.133 | 0.123 |

| | | | | | | | | | | | |
|---|---|---|---|---|---|---|---|---|---|---|---|
| | | 26 | F7 | 24-Dec | 25-Dec | 26-Dec | petite | 0.095 | -0.076 | 0.122 | 0.080 |
| | | 27 | B9 | 26-Dec | 26-Dec | 30-Dec | grande | 0.133 | -0.088 | 0.119 | 0.120 |
| | | 28 | F8 | 24-Dec | 26-Dec | 30-Dec | petite | 0.133 | -0.095 | 0.133 | 0.086 |
| Trial 1 13-Dec | NiZn | 01 | B7 | 21-Dec | 24-Dec | 26-Dec | grande | 0.053 | -0.045 | 0.056 | -0.024 |
| | | 02 | F5 | 21-Dec | 24-Dec | 26-Dec | grande | 0.063 | -0.030 | 0.057 | -0.033 |
| | | 03 | D6 | 23-Dec | 25-Dec | 29-Dec | grande | 0.058 | -0.037 | 0.050 | -0.030 |
| | | 04 | B2 | 26-Dec | 26-Dec | 30-Dec | grande | 0.047 | -0.052 | 0.053 | -0.037 |
| | | 05 | D8 | 23-Dec | 26-Dec | 30-Dec | grande | 0.058 | -0.039 | 0.046 | -0.037 |
| | | 06 | E4 | 26-Dec | 26-Dec | 30-Dec | petite | 0.026 | -0.123 | -0.022 | -0.051 |
| | | 07 | H9 | 26-Dec | 26-Dec | 30-Dec | grande | 0.021 | 0.006 | 0.022 | -0.006 |
| Trial 2 23-May | CdMn | 01 | F6 | 26-May | 26-May | 28-May | petite | 0.096 | -0.074 | 0.085 | 0.021 |
| | | 02 | G7 | 26-May | 26-May | 30-May | petite | -0.019 | -0.107 | 0.084 | -0.081 |
| | | 03 | H9 | 26-May | 26-May | 28-May | petite | 0.020 | -0.083 | 0.105 | -0.092 |
| | | 04 | B4 | 28-May | 28-May | 30-May | petite | 0.073 | -0.104 | 0.101 | 0.043 |
| | | 05 | C3 | 28-May | 28-May | 30-May | petite | 0.018 | -0.072 | 0.110 | -0.036 |
| | | 06 | D8 | 28-May | 28-May | 30-May | petite | 0.020 | -0.079 | 0.105 | 0.011 |
| | | 07 | B3 | 31-May | 31-May | 04-Jun | petite | 0.020 | -0.079 | 0.114 | -0.026 |
| | | 08 | E9 | 29-May | 31-May | 03-Jun | petite | 0.025 | -0.161 | 0.022 | -0.045 |
| | | 09 | C7 | 31-May | 02-Jun | 05-Jun | grande | 0.042 | -0.091 | 0.055 | 0.001 |
| | | 10 | F8 | 31-May | 02-Jun | 04-Jun | petite | 0.084 | -0.115 | 0.070 | -0.029 |
| | | 11 | E7 | 05-Jun | 05-Jun | 07-Jun | petite | -0.003 | -0.105 | 0.083 | -0.048 |
| Trial 2 23-May | CoCu | 01 | B10 | 28-May | 28-May | 01-Jun | petite | 0.053 | -0.108 | -0.069 | -0.085 |
| | | 02 | E3 | 28-May | 28-May | 01-Jun | petite | 0.099 | -0.085 | -0.068 | -0.076 |
| | | 03 | F9 | 26-May | 28-May | 31-May | petite | 0.057 | -0.150 | -0.123 | -0.107 |
| | | 04 | F11 | 26-May | 28-May | 06-Jun | NA | NA | NA | NA | NA |
| | | 05 | G11 | 26-May | 28-May | 01-Jun | petite | 0.100 | -0.114 | -0.080 | -0.066 |
| | | 06 | H10 | 28-May | 28-May | 31-May | petite | 0.094 | -0.099 | -0.072 | -0.073 |
| | | 07 | A9 | 28-May | 29-May | 31-May | petite | 0.031 | -0.142 | -0.140 | -0.135 |
| | | 08 | G10 | 28-May | 29-May | 31-May | petite | 0.088 | -0.089 | -0.068 | -0.091 |
| | | 09 | H11 | 29-May | 29-May | 02-Jun | petite | 0.082 | -0.094 | -0.063 | -0.084 |
| | | 10 | A8 | 28-May | 31-May | 02-Jun | petite | 0.086 | -0.120 | -0.062 | -0.071 |
| | | 11 | F8 | 28-May | 31-May | 04-Jun | petite | 0.012 | -0.138 | -0.115 | -0.127 |
| | | 12 | H4 | 28-May | 31-May | 04-Jun | petite | 0.066 | -0.096 | -0.089 | -0.080 |
| | | 13 | A7 | 02-Jun | 02-Jun | 07-Jun | NA | NA | NA | NA | NA |
| | | 14 | A11 | 02-Jun | 02-Jun | 04-Jun | petite | 0.108 | -0.115 | -0.062 | -0.083 |
| | | 15 | A5 | 02-Jun | 03-Jun | 06-Jun | petite | 0.053 | -0.129 | -0.095 | -0.105 |
| | | 16 | A6 | 06-Jun | 06-Jun | 07-Jun | petite | 0.087 | -0.087 | -0.070 | -0.047 |
| | | 17 | A10 | 06-Jun | 06-Jun | 07-Jun | petite | 0.042 | -0.099 | -0.120 | -0.113 |

Table 4: Fitness assays: Information on fitness assays including date of experiment, combos tested; negative controls as maximum growth rates calculated from noise in blank media (no yeast added; average from 3 technical replicates in metal environments and 4 to 30 technical replicates in YPAD); max. growth rates of the ancestor in all test environments (average of three technical replicates in metal environments and 4 to 33 technical replicates in YPAD); and average relative fitnesses of evolved lines in each test environment.

| | Batch | combofit_1 | combofit_2 | combofit_3 | | combofit_4 | combofit_5 | combofit_6 | combofit_7 |
|---|---|---|---|---|---|---|---|---|---|
| | Date | 28-Aug-24 | 06-Sep-24 | 17-Sep-24 | | 23-Sep-24 | 26-Sep-24 | 01-Oct-24 | 09-Jan-25 |
| | Metal combination | CoMn | CoCu | CdMn | NiZn | MnNi | CdCu | CdNi | CuMn |
| Negative controls | $r_{YPAD}^{Blank}$ | 0.014 | 0.011 | 0.016 | 0.016 | 0.020 | 0.015 | 0.013 | 0.016 |
| | $r_{Combo}^{Blank}$ | 0.018 | 0.006 | 0.024 | 0.017 | 0.011 | 0.006 | 0.011 | 0.011 |
| | $r_{S}^{Blank}$ | 0.018 | 0.004 | 0.015 | 0.024 | 0.037 | 0.007 | 0.004 | 0.004 |
| | $r_{W}^{Blank}$ | 0.008 | 0.008 | 0.023 | 0.022 | 0.023 | 0.006 | 0.015 | 0.012 |
| Ancestral growth rates | $r_{YPAD}^{Anc}$ | 0.281 | 0.310 | 0.295 | 0.295 | 0.301 | 0.309 | 0.320 | 0.301 |
| | $r_{Combo}^{Anc}$ | 0.053 | 0.046 | 0.074 | 0.070 | 0.059 | 0.113 | 0.023 | 0.072 |
| | $r_{S}^{Anc}$ | 0.119 | 0.257 | 0.107 | 0.192 | 0.077 | 0.205 | 0.082 | 0.057 |
| | $r_{W}^{Anc}$ | 0.095 | 0.287 | 0.130 | 0.227 | 0.100 | 0.196 | 0.061 | 0.044 |
| Avg. relative fitness $(r_{Evo} - r_{Anc})$ | $\bar{f}_{YPAD}$ | -0.112 | -0.111 | -0.097 | -0.046 | -0.081 | -0.046 | -0.160 | -0.095 |
| | $\bar{f}_{Combo}$ | 0.090 | 0.070 | 0.034 | 0.046 | 0.114 | 0.014 | 0.069 | 0.043 |
| | $\bar{f}_{S}$ | 0.048 | -0.086 | 0.085 | 0.037 | 0.127 | -0.013 | 0.073 | 0.119 |
| | $\bar{f}_{W}$ | 0.051 | -0.089 | -0.025 | -0.031 | 0.096 | -0.055 | 0.000 | 0.044 |

### Supplementary Figures

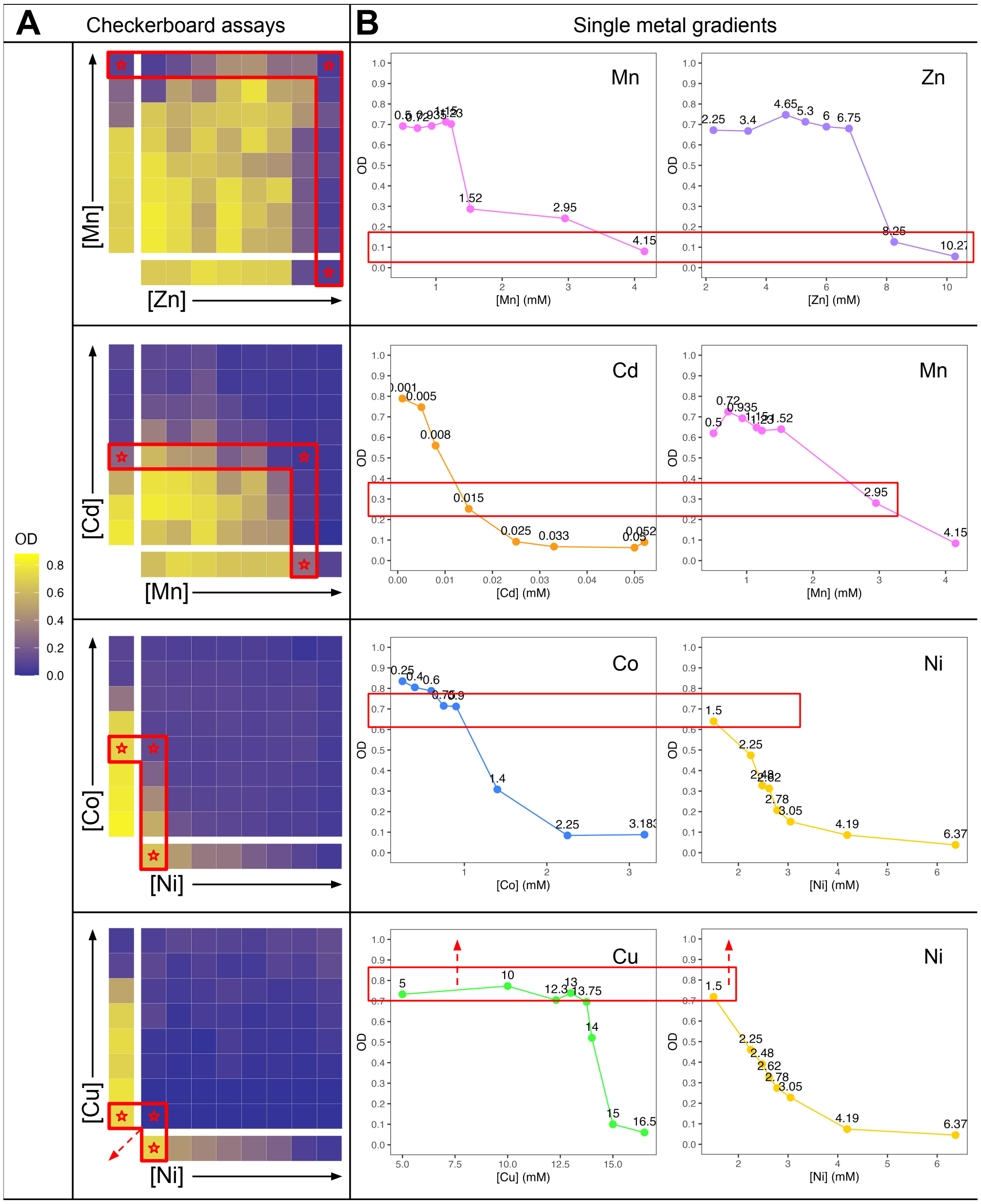

Figure 1: The first row shows results from MnZn ($a$ = 2.68, qualitative category 1), the second row is CdMn ($a$ = 1.25, qualitative category 2), the third row is CoNi ($a$ = 0.82, qualitative category 3), and the fourth row is CuNi ($a$ = 0.98, qualitative category 4) (A) Checkerboard assay results. Color in the heatmap represents OD48hr ranging from ∼0.0 to ∼0.9 after subtraction of absorbance reading from blank media (legend on left). Red stars represent concentrations that meet the two goals; red stars on x and y axes denote single metal concentrations that produce equivalent OD, and red star in the central grid shows that, when combined, these concentrations produce an OD near 0. (B) Numerical values of single metal OD48hr and concentrations. The graphs on the left correspond to the y axes in panel A, and the graphs on the right correspond to the x axes. Metal identity is shown in the top right. Red boxes overlay the ODs produced by the red star concentrations in panel A. Red arrows in the last row, CuNi, show that the concentrations may need to be further decreased since growth was not observed at any concentration tested.
